# Goat kid recognition of their mothers’ calls is not impacted by changes in source-filter parameters

**DOI:** 10.1101/2022.05.19.492593

**Authors:** Tania Perroux, Alan G. McElligott, Elodie F. Briefer

## Abstract

Features varying more between than within individuals are usually considered as potential cues for individual recognition. According to the source-filter theory of vocal production, the fundamental frequency of mammal’s vocalisations depends on the characteristics of the vocal folds, while formants are determined by the characteristics of the vocal tract. Goat mothers and their kids (*Capra hircus*) display mutual recognition, and both source-related parameters (F0) and filter-related ones (formants) have been shown to be individualised in their vocalisations. Here, we aimed to identify if these parameters (source-related parameters (F0) and/or filter vocal parameters) are used by goat kids to recognise their mother’s vocalisations. To this aim, we used an algorithm to modify either F0 or formants of the calls of goat mothers to different degrees (within or exceeding the range of natural intra-individual variability), and we played back these modified calls to their kids. We did not observe any difference in the kid reactions to the modified maternal vocalisations and to the natural calls. We suggest that either: *(i)* fundamental frequency and formants are not involved in maternal recognition in goats; *(ii)* goat kids have a tolerance for variation when recognising their mother’s calls that exceeds the shifts we performed; *(iii)* goat maternal recognition is based on other vocal features than those tested here, or (*iv*) goat kid maternal recognition is based on a combination of different features and might be more flexible than previously thought, such that when one main feature is modified, kids focus on other features.

## Introduction

Individual recognition is crucial for directed parental care (Briefer & McElligott, 2011a; Gokcekus *et al*., 2021; Searby & Jouventin, 2003), as well as for offspring survival (Padilla de la Torre *et al*., 2016). Parent-offspring recognition develops quickly and is influenced by environmental constraints (Briefer *et al*., 2012). Mother-offspring recognition at a distance mostly relies on visual and acoustic cues to infer the position of the young/parent, while recognition at close quarters is mostly sustained by olfaction (Ferreira *et al*., 2000; Torriani, Vannoni & McElligott, 2006). In larger groups where risks of confusion are enhanced, accurate parent-offspring recognition prevents misdirected maternal care; particularly when the neonate depends entirely on its mother for food and/or when lactation requires a lot of energy (Sèbe *et al*., 2010; Linossier *et al*., 2021).

Vocalisations play an important role in individual recognition (Yorzinski, 2017). To enable vocal recognition, features typically vary more between than within individuals (Li *et al*., 2017). In species where contact calls have been shown to be individualised, offspring react more to the calls of their mothers compared to calls from other females. For example in ungulates, calves *(Bos taurus)* stay for longer near and approach closer a loudspeaker broadcasting their mother’s voice compared to one broadcasting another female’s voice (Padilla de la Torre *et al*., 2016). Ungulate offspring can use cues in both the frequency and temporal domains for maternal recognition (Charrier, Mathevon & Jouventin, 2003), with for instance sheep (*Ovis aries*) showing an early filial preference that depends on acoustic cues (Sèbe *et al*., 2007, 2010).

In goats (*Capra hircus)*, mothers and kids have individualised contact calls (Briefer & McElligott, 2011a). Goats vocalisations are characterised by a clear harmonic structure, as well as strong frequency and amplitude modulations (Briefer *et al*., 2012). When isolated, goats produce two types of calls: contact calls and isolation calls, the latter characterised by higher pitch (Briefer & McElligott, 2011a; Siebert *et al*., 2011). Contact calls are produced either open or closed mouthed, with close mouth altering the formant structure (Favaro, Briefer & McElligott, 2014). Sibling goat kids have more similar vocalisations compared to unrelated goat kids, and this effect is not dependent on sex nor experience (Burke *et al*., 2020). However, goats also show some flexibility in their calls during development: the call of young kids living in the same group converge over time and become more similar than those raised in different groups (Briefer & McElligott, 2012).

Goat mother-offspring relationships are characterised by a specific, rapidly formed and fairly stable maternal attachment (Hernández *et al*., 2012), leading to mutual recognition (Briefer & McElligott, 2011a). Mothers develop acoustic recognition and discrimination of their kid’s contact call, which remains even after weaning and long-term separation (Briefer *et al*., 2012). Similarly, from five days old at least, goat kids have been shown to differentiate between the calls of their mothers and other familiar females based on vocal cues (Briefer & McElligott, 2011a), although the precise features used in vocal recognition are not yet known. This mutual mother-kid recognition could be based notably on the fundamental frequency (F0; “source”, which depends on the characteristics of the vocal fold) and/or on the formants (“filter”, which are determined by the characteristics of the vocal tract). Indeed, both F0 and formants have previously been identified as potential markers of individuality in goat kids’ bleats (Briefer & McElligott, 2011a).

There are two types of paradigms to evaluate the cues used in vocal discrimination. First, investigating the extent to which vocal features are stable within individuals (*i*.*e*., stereotypic) can be used to determine the likelihood that a certain feature will be involved in individual recognition (Pitcher, Harcourt & Charrier, 2012; Sauvé *et al*., 2015). Second, playbacks can be carried out with modified calls in which features potentially used for individual recognition are altered one by one, while keeping the rest constant. These modified calls can then be played to the animals to assess whether these changes impair recognition (Charrier *et al*., 2003; Tamura *et al*., 2021). If recognition is impaired by the modification of a parameter, this suggests that it is used for recognition. Such experiments have shown, for example, that Australian sea lions (*Neophoca cinerea*) pups respond less to their mother’s calls when F0 has been shifted (once, twice or three times the standard deviation) than to natural calls, suggesting that they use this vocal feature for maternal recognition (Charrier, Pitcher & Harcourt, 2009).

The present study was focus on determining if source-filter vocal parameters (*e*.*g*., F0 and/or formant frequencies) are used by goat kids to recognise their mother’s vocalisations. To this aim, we played back the vocalisations of mothers to their kids, where either F0 or formants had been modified to different degrees (within or exceeding the range of natural intra-individual variability). We predicted that, if a feature is involved in vocal recognition, kids would react less to the modified vocalisation than to the natural one, as the modification impairs recognition (Charlton, Huang & Swaisgood, 2009a; Charrier *et al*., 2009).

## Material and Methods

### Subjects and housing

Our subjects were 14 goat kids (six females and eight males), born from seven multiparous pygmy goat mothers and the same father in spring (n = 10) and summer 2011 (n = 4). All kids were born and raised at the WhitePost Farm (53°06’N, 1°03’W, UK). Kids’ age ranged from 10 to 28 days (mean 17.08 ± 5.28 days) at the time of playbacks. The goats used in this study were kept indoors in a communal pen of 4.4 m × 4.5 m. Following the husbandry routine carried out by the farm employees, females who were about to give birth were isolated in a 2.5 m^2^ pen within the communal pen and kept there with their kid(s) for two to three days. The aim was to allow adequate development of the mother–offspring relationship and prevent interference from other goats. Mothers and kids were then released in the communal pen.

### Playback preparation

The mothers’ calls were recorded two to five days before the playbacks, by separating kids from their mothers behind a fence (1–10 m) for no more than five minutes, two times a day, to elicit contact calls. Calls were recorded at a distance of 1–5 m from the mother using a Sennheiser MKH70 connected to a Marantz PMD660 recorder (sampling rate: 44.1 kHz) (see Briefer and McElligott, 2011b, 2012 for further details about the procedure). Open-mouth contact calls were then saved on a computer in wav, 16-bit, and visualised on spectrograms in Praat v.5.0.47 DSP Package (Boersma and Weenink, 2009) (window length = 0.01 s, dynamic range = 50 dB). Eight good-quality calls per individual (low level of background noise) were selected for preparing the playback treatments.

To determine to which degree F0 and formants should be modified, the intra and inter-individual variability of the mean F0 frequency value across the call (‘F0Mean’) and the mean frequency value of the fourth formant (‘F4Mean’) were measured based on eight calls from 11 adult goats recorded previously (Briefer & McElligott, 2011b). This sample of individuals included the seven mother goats whose calls were played back to kids in the current study. The fourth formant was chosen for analyses as it is the most salient and easy to measure (Briefer & McElligott, 2011b). The intra-individual variation (maximum – minimum value for each individual) was as follows (mean ± SD): F0Mean = 66.05 ± 36.52 Hz, F4Mean = 521.24 ± 262.17 Hz. The first modified call treatment was aimed to mimic a shift in F0 or formants that was within the extreme range of within-individual variation, while the second treatment was aimed to mimic a shift outside this range. Based on the intra-individual variation values, we determined the first shift to be of about 70 Hz above the natural signal for F0Mean (‘F0 Shift1’) and of about 520 Hz for F4Mean (‘Formant Shift1’), and of about twice these values for the second shift (160 Hz for F0Mean (‘F0Shift2’) and 1040 Hz for F4Mean (‘Formant Shift2’)).

The preparation and modification of the sequences to play back was carried out in Praat as follows: the eight selected calls were inserted in a sequence, interspaced by intervals of natural duration (0.98 s for adult goats: Briefer & McElligott, 2011b), made of the goat’s usual background noise. The sequence was then repeated to obtain a 30 s long sequence. For twin kids (n = 7 pairs), the same mother calls were used but they were inserted in a different order in the sequence. All calls in a given sequence were then rescaled to the same maximum amplitude. Following this sequence preparation, F0 and formants were modified independently using a PSOLA-based algorithm with a custom Praat script (Pitcher, Briefer & McElligott, 2015), which shifted the F0 or formant up by a predetermined resynthesis factor, supposedly leaving the other acoustic parameters (*e*.*g*., formant and F0, respectively) unchanged. To define the predetermined resynthesis factor needed, the first call of each sequence was analysed to extract its F0Mean and F4Mean. Based on these extracted values, a resynthesis factor of 1.21 to 1.80 was used for F0 Shift1, and of 1.50 to 2.82 for F0 Shift2. For F4Mean, a resynthesis factor of 0.85 to 0.89 was used for Formant Shift1, and of 0.74 to 0.80 for Formant Shift2.

Modified calls of the mothers were examined to verify the modification and the absence of artefacts (Fig. 1). In addition, the eight vocalisations constituting the sequence for each mother and in each treatment were then analysed using a script adapted from Reby and McComb (2003) and Charlton, Zhihe & Snyder (2009b), to obtain the precise F0 and formants values of all natural and modified calls played back. To do so, the following settings were used: Source-related vocal parameters (F0 mean, minimum, maximum values and range) were measured by extracting the fundamental frequency contour of each call using a cross-correlation method ([Call: To Pitch (cc) command], F0: time step = 0.01 s, pitch floor = 70 Hz, pitch ceiling = 750 Hz). Filter-related (formants) vocal parameters (F1, F2, F3 and F4 mean values and formant dispersion) were measured by extracting the contour of the first four formants of each call using Linear Predictive Coding analysis (LPC; [Call: To Formant (burg) command], time step = 0.01 s, maximum number of formants = 4, maximum formant = 5000 Hz (natural calls) to 6750 Hz (Formant Shift2), window length = 0.1 s).

**Figure 1:**
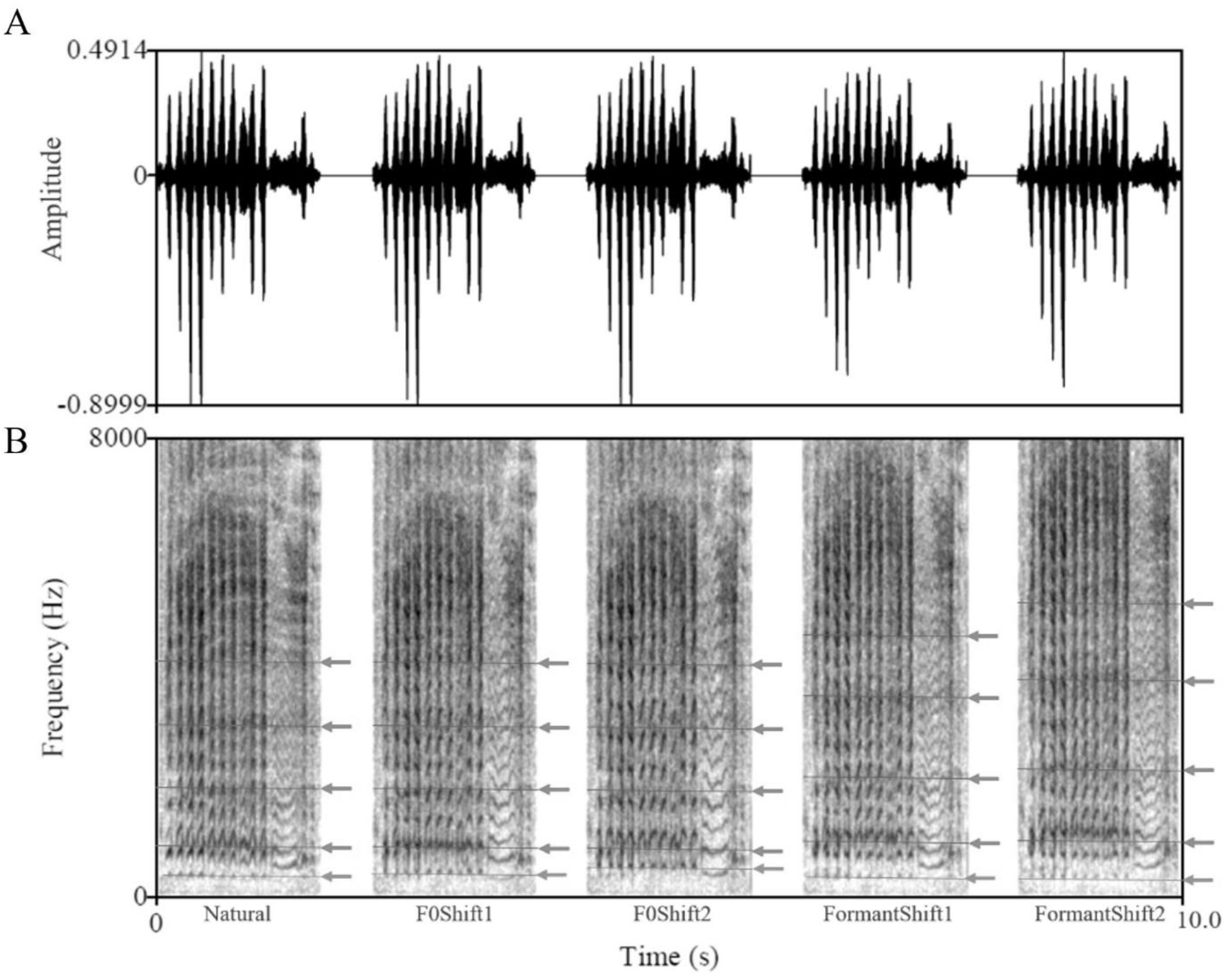
Example of a mother goat call oscillogram (A) and spectrogram (B) for each treatment. Arrows on the spectrograms indicate the Fundamental frequency F0 (blue arrow) and the first four formants (red arrow).

### Experimental set up

The playback experiment was carried out in a 2.5 m^2^ arena situated within the same barn but outside visual and hearing range from the kids’ home pen. The testing arena was placed within a pen containing other species, to which the subjects were habituated to (*i*.*e*., sheep and llamas), surrounded on two sides by concrete walls, and on the two others by goat fences (Fig. 2). To prevent disturbance from other animals, one side was covered with a blanket. The loudspeaker and the camera were placed on the adjacent side, at about 2-4 m from the subject. The floor was covered with straw. Subjects were habituated to the pen for five minutes per day, alone, during three to four days before the first playback trial started.

**Figure 2:**
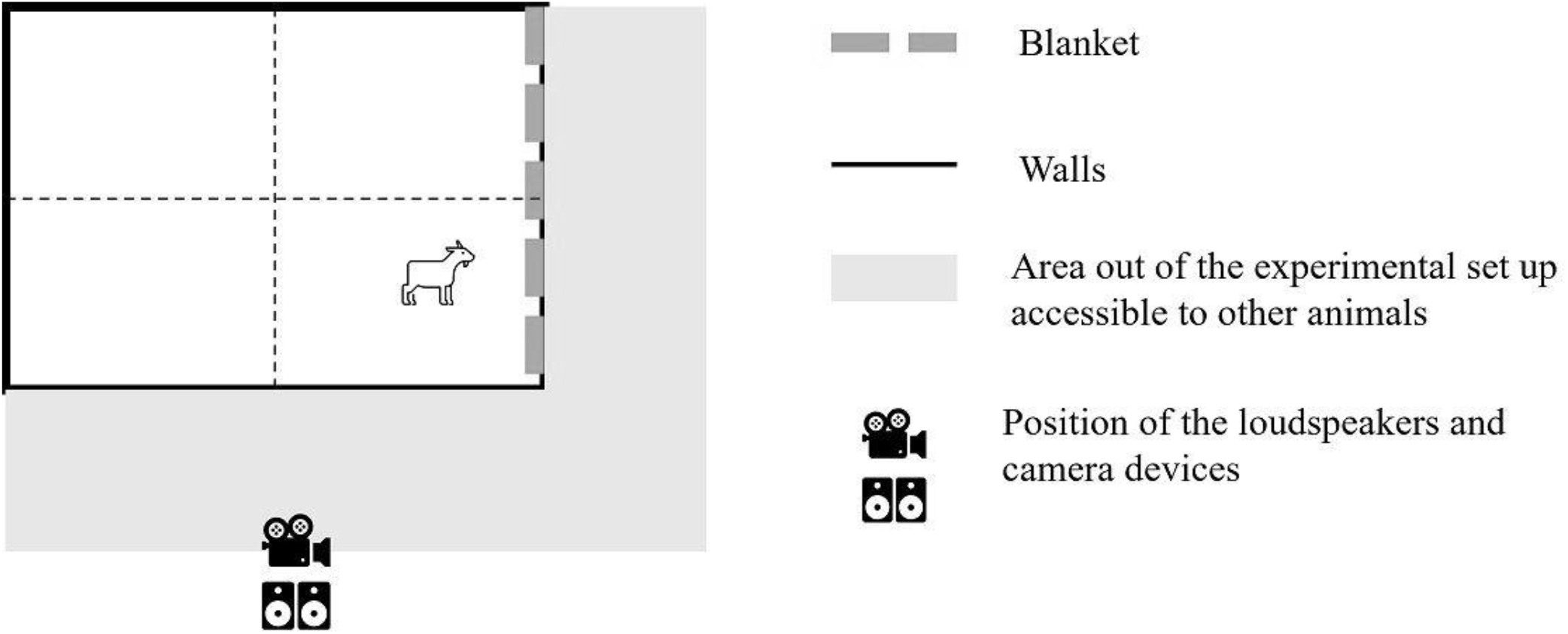
Schematic representation of the experimental set used during the playback sessions. Loudspeaker and camera devices were positioned about two meters away from the fences. A blanket was installed to reduce disturbance from surrounding animals that could pass by the experimental set up.

### Experimental procedure

Following Briefer and McElligott (2011b), after ensuring that it suckled from its mother before starting the procedure, each kid was placed with other pen mates (two to four kids together) for 1.5 to 2 h in the experimental set up before the first trial started, to trigger responses to maternal calls and allow the subjects to habituate to the pen. At the end of this habituation period, the other pen mates were removed and the playback experiment started when subject was settled (*i*.*e*., did not show any obvious signs of stress, such as calling or defecating). The call treatments, stored as high-quality mp3 files (sampling rate = 44.1 kHz; bit rate = 224 kbps), were played using a Skytronic TEC076 portable system (frequency response: 50 Hz– 20 kHz ± 3 dB), at an intensity estimated to be normal for the goats (80 dB at 1 m; Briefer & McElligott, 2011b). Each kid was tested with the five playback treatments on two to three consecutive days (one to three treatments per day). To ensure relatively similar conditions and kid age between playbacks of F0 and formant modifications, respectively, the order of testing condition were pseudo-randomised: the natural condition could take place in any of the five trials, but F0 tests would take place on either the first, second or third trial of testing whereas formant tests would occur on either the third, fourth or fifth trial. The behavioural response of the animals was recorded using a Sony DCR-SX50E camcorder. Kids were returned to their mother directly after each test.

### Video analysis

The videos of the playbacks were scored while blind to the treatment. The behaviours were scored continuously, during the period preceding the playback (‘Pre-playback’; duration: 40.93 s ± 6.06) and the rest of the video corresponding to the playback itself (‘Playback’; duration: 37.27 s ± 4.37) using the software BORIS v7.9.8 (Friard & Gamba, 2016). The coded behaviours were as follow: looking towards the loudspeaker (evaluated with a 45° angle of head in direction to the device), locomotion (with four legs moving), call (vocalisations of the goat kid) and latencies of the first call, locomotion and look towards the speaker after the playback onset. Except for latencies, all behaviours were divided by the duration of the experimental phase for further analysis, resulting in ‘rates’ for events (number of occurrence / observation duration), and ‘ratios’ for states (duration of the state / observation duration).

To ensure reliability of the video coding, the intra-observer reliability (Bateson & Martin, 2021) was calculated using the following procedure: 10 randomly selected videos were encoded twice in a random order. For each behaviour, a correlation coefficient (r^2^) was then calculated using a Pearson correlation test. For all behaviours, we obtained an r^2^ ≥ 0.79 (mean = 0.95; range = 0.79-0.99), suggesting good intra-observer reliability *(Table S1)*.

### Statistical Analysis

Statistical analyses were conducted with RStudio (v1.3, R Core Team). The effects of F0 and formant manipulations were analysed separately. The dataset was hence split as follows: F0 conditions (natural, F0 shift1 and F0 Shift2) and Formant conditions (natural, Formant Shift1 and Formant Shift2). The same procedure (points 1-3 below) was applied to the two sub-datasets.

1. For behaviours that could be measured before and after the playback onset (*i*.*e*., all except the latencies to respond to the playback), a comparison was made between these two periods to select behaviours that could be considered as a reaction to the calls (*i*.*e*., that were affected by the auditory stimuli). To this aim, a Linear Mixed-Effect Models (LMM) was ran for each behavioural response, including the behavioural response as the outcome variable, the period (before or after playback) as a fixed factor and the identity of the subject nested within the identity of its mother as a random factor. Behaviours that differed significantly between the two scoring periods were selected for further analyses.
2. To test for the effect of the playback treatment on the responses of the kids, for each extracted acoustic parameter, the mean acoustic values of the calls of each mother in each treatment were subtracted to the mean value of the natural treatment, to obtain the actual shift (*i*.*e*., the natural vocalisation was therefore fixed at zero, and each value given actually refers to that difference, hereafter “playback shift”). One LMM model was built for each selected behaviour, entered as an outcome variable, and for each acoustic parameter measured (F0: mean (‘Mean F0’), minimum (‘Min F0’), maximum (‘Max F0’) values and range (‘Range F0’); and formants: (Mean F1, F2, F3 and F4 values and formant dispersion), whose playback shift values were entered as a continuous fixed factor (Total: 24 models for the F0 data subset, and 30 for the formant subset). In addition, we included the order of the treatments (one to five), as well as the pre-playback behaviour, as fixed continuous control factors. The identity of the subject nested within the identity of its mother was used as a random effect to control for repeated measurements of the same subjects and potential similarities between twins.
3. We then investigated the impact of modifying F0 on formants, and vice versa, in order to validate our PSOLA-algorithm procedure. To this aim, we ran further LMMs including the mean value, for each playback sequence of each mother’s calls, of the vocal parameters extracted from the acoustic analyses of the calls played back as an outcome variable (in separate models, F0 values: Mean F0, Min F0, Max F0 and Range F0; and formants values: Mean F1, F2, F3 and F4 values and formant dispersion). Each model included the playback condition as a fixed effect (*i*.*e*., Natural, Formant Shift1 and Formant Shift2 for LMM carried out on F0 values and Natural, F0 Shift1 and F0 Shift2 for LMM carried out on formant values), and the identity of the mother as a random factor. For models where the treatment had an effect on the acoustic features, post-hoc Tukey tests were conducted. The results of these models can be found in supplementary material (*Tables S2 and S3*).

For all LMMs, we checked the residuals of the models graphically for normal distribution and homoscedasticity (simulateResiduals function, package DHARMa, Hartig, 2022). If the assumptions were not met, a logarithmic transformation was used. When the assumptions of normality and homoscedasticity were not met despite a logarithmic transformation, the data were transformed to binary data (superior to the median in the treatment = 1, inferior to the median = 0) and input into Generalized Linear Mixed-Effect Models (GLMM) instead of a LMM, with the same fixed, control and random factors (function glmer, package lme4; Bates *et al*., 2015). Precise types of models can be found in the supplementary material (*Table S4*). P-values were calculated by comparing model with and a model without the term of interest using parametric bootstrap methods (1000 bootstrap samples; PBmodcomp function, package pbkrtest, Halekoh & Højsgaard, 2014). To this aim, models were fitted with maximum likelihood.

One of the mothers died due to causes unrelated to the experiment during the testing period (between playback sessions). To ensure that the responses of her twin offspring did not differ from those of other kids, Wilcoxon signed rank exact tests were used to compare the mean values of each behaviour in each treatment including the responses from these two kids with the variance when the response of these kids after their mother’s death were excluded (three trials for one kid and four trials for the other kid). Since these differences were not significant (F0: V = 39, p-value = 0.144; Formants: V = 88, p-value = 0.546), the responses of these two kids were included in analyses.

### Ethical Note

Animal care and all experimental procedures were in accordance with the International Society for Applied Ethology guidelines(“Guidelines for the treatment of animals in behavioural research and teaching,” 2012). The experiments were carried out in 2011. At that time, no ethical approval was required in the UK for such non-invasive playback experiments.

## Results

### Behaviours affected by the playback

For both F0 and formant conditions, the following goat kid behaviours significantly differed between before and after the playback onset: call rate (LMM: F0, p<0.0001; formants, p<0.0001), locomotion ratio (LMM: F0, p<0.05; formants, p<0.05) and looking ratio (LMM: F0, p<0.01; formants, p<0.01). These behaviours and the corresponding latencies were therefore selected for further analysis.

### Effect of F0 modifications on maternal recognition

F0 modifications did not affect kid responses to the playbacks (Table 1; *e*.*g*., Fig. 3). The order in which goat kids underwent the treatments had a significant effect on nine models for behaviours coded as rates and ratios (call, locomotion and looking towards the speaker) out of 12 and did not impacted the models for latencies *(Table S5)*. Goat kid’s behaviour before playback onset affected their behaviour after the onset on all models for all behaviours coded as rates and ratios *(Table S6)*.

**Table 1:** Effect of actual shifts in F0 among playbacks of natural calls (no shift), F0 Shift1 (*i*.*e*., within the natural variability) and F0 Shift2 (*i*.*e*., exceeding the natural variability of the mother’s vocalisation) on goat kids’ behaviours. F0 values were obtained by subtracting each mean frequency value of the playback sequence to the mean values of the natural playback of the corresponding individual, giving a value of zero for the natural call. Linear Mixed-Effect Models (LMMs; p values extracted using parametric bootstrap) did not reveal any significant effect of the shifts on goat kids’ behaviour. In bold are the lowest p values, whose relationships are illustrated in Fig. 3

| P-values | Latency<br>Call | Latency<br>Locomotion | Latency<br>Look | Call Rate | Locomotion<br>Ratio | Looking<br>Ratio |
| --- | --- | --- | --- | --- | --- | --- |
| Max F0 | 0.652 | 0.377 | 0.366 | 0.649 | 0.289 | 0.646 |
| Mean F0 | 0.519 | 0.293 | 0.247 | 0.377 | 0.159 | 0.433 |
| Min F0 | 0.484 | 0.515 | 0.236 | 0.482 | 0.156 | 0.291 |
| Range F0 | 0.851 | 0.624 | 0.989 | 0.738 | 0.766 | 0.442 |

**Table 2:** Effect of actual shifts in Formants among playbacks of natural calls (no shift), Formant Shift1 (i.e., within the natural variability) and Formant Shift2 (i.e., exceeding the natural variability of the mother’s vocalisation) on goat kids’ behaviours. Formant values were obtained by subtracting each mean frequency value of the playback sequence to the mean values of the natural playback of the corresponding individual, giving a value of zero for the natural call. Linear Mixed-Effect Models (LMMs; p values extracted using parametric bootstrap) found no significant effect of the shifts on goat kids’ behaviour. In bold are the lowest p values, whose relationships are shown in Fig. 4.

| P-values | Latency<br>Call | Latency<br>Locomotion | Latency<br>Look | Call<br>Rate | Locomotion<br>Ratio | Looking<br>Ratio |
| --- | --- | --- | --- | --- | --- | --- |
| Mean F1 | 0.306 | 0.506 | 0.43 | 0.934 | 0.402 | 0.69 |
| Mean F2 | 0.400 | 0.453 | 0.715 | 0.924 | 0.470 | 0.727 |
| Mean F3 | 0.406 | 0.573 | 0.526 | 0.923 | 0.450 | 0.796 |
| Mean F4 | 0.826 | 0.566 | 0.909 | 0.757 | 0.125 | 0.364 |
| Dispersion | 0.545 | 0.156 | 0.744 | 0.687 | 0.128 | 0.336 |

**Figure 3:**
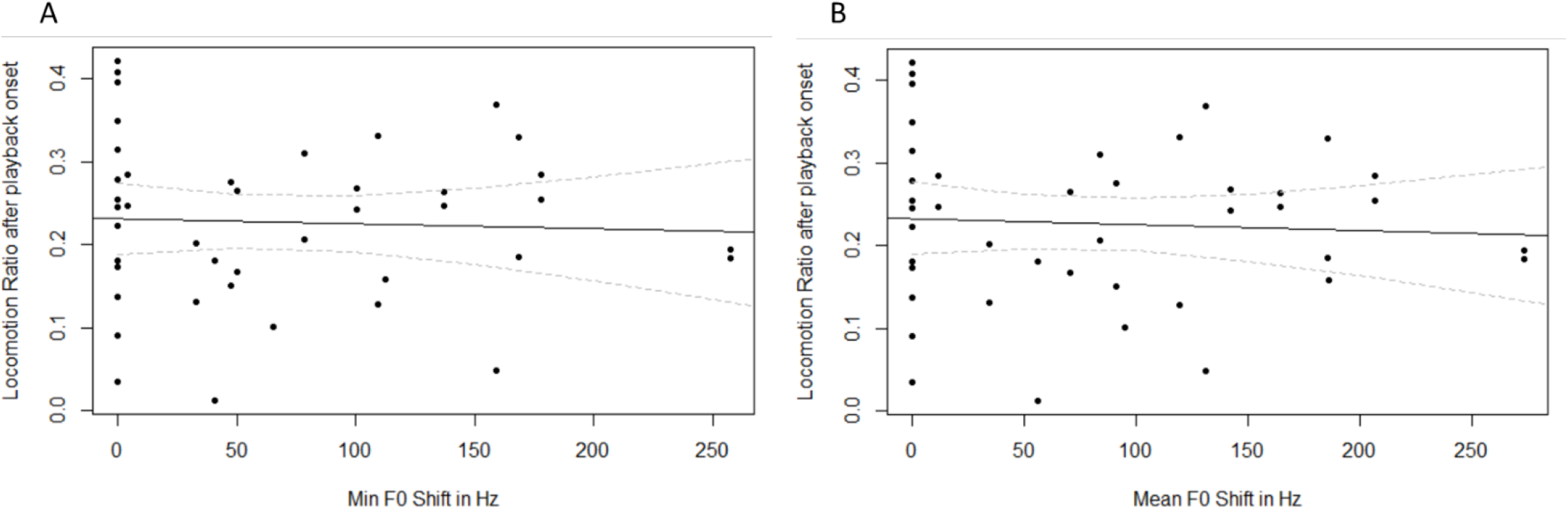
Effect of the actual shift in minimum (A) and mean (B) F0 on locomotion ratio (time spent performing the behaviour divided by the total duration of the video sequence) after playback onset over the three treatments: natural voice of the mother (set at 0 Hz), a positive shift within the natural range (about 70 HZ above natural F0 Mean) and a positive shift exceeding the natural range (about 160 HZ above natural F0 Mean). Dots represent the data, and the black line represent the predicted effect given by the LMM model. In grey are given the 95% predicted confidence intervals.

### Effect of formants modifications on maternal recognition

Formant modifications did not affect kid responses to the playbacks (Table 4; *e*.*g*., Fig. 4). The order in which kids underwent the treatments had a significant effect on all models for locomotion and looking towards the loudspeaker latencies, and for call and locomotion ratios *(Table S7)*. Goat kid’s behaviour before playback onset affected their behaviour after the onset on all models for call rate and locomotion ratio *(Table S8)*.

**Figure 4:**
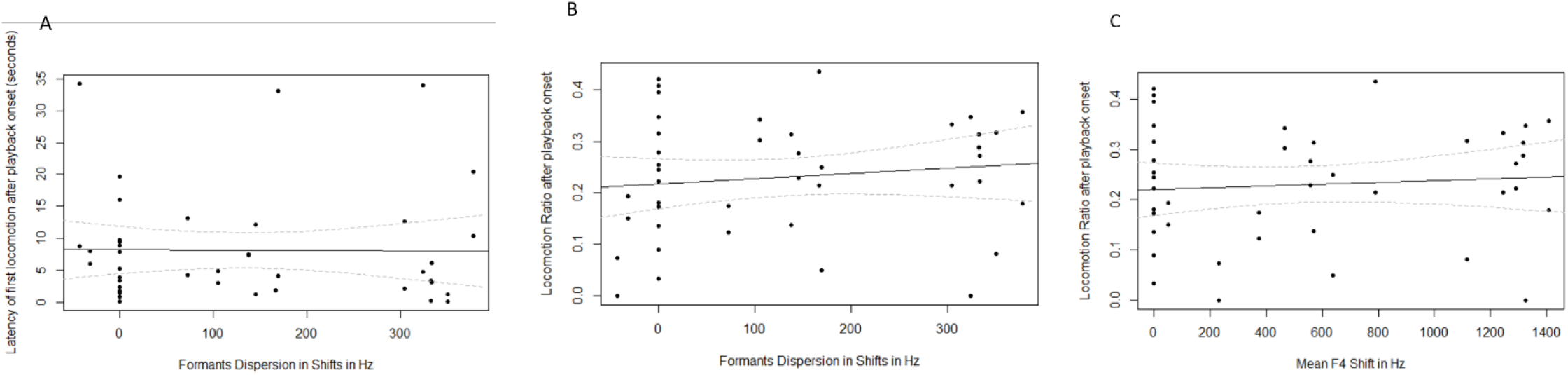
Effect of the actual shift in formant dispersion on latency of the first locomotion (A) and locomotion ratio (B), and of the shift in mean value of F4 on locomotion ratio (C) after playback onset over the three conditions: natural voice of the mother (set at 0 Hz), a positive shift within the natural range (about 520 HZ above natural F4 Mean), and a positive shift exceeding the natural range (about 1040 HZ above natural F4 Mean). Ratios were obtained by dividing the duration of the behaviour by the total duration of the video sequence. Dots represent the data, and the black line represent the predicted effect given by the LMM model. In grey are given the 95% predicted confidence intervals.

## Discussion

Mother goats and their kids display mutual recognition, and kids recognise their mother based on vocal cues from at least five days old (Briefer & McElligott, 2011a). However, the vocal parameters used for achieving this vocal recognition remain unknown. We investigated whether goat kids would react to modified versions of their mother’s calls, using two types of changes (to fundamental frequency and formants) and two intensities of modifications (within the intra-individual variability or exceeding this variability). We found that call rate, locomotion ratio and looking ratio were affected by the playback onset. However, goat kids responded as much when exposed to natural maternal vocalisations, compared to when the fundamental frequency or the formants were modified. We suggest that goat kids recognise their mother vocalisations based on several possible non-exclusive mechanisms: (*i)* fundamental frequency and formants are not involved in maternal recognition in goats (Carlson, Kelly & Couzin, 2020); *(ii)* goat kid maternal recognition sensitivity exceeds the shifts we performed (Aubin & Jouventin, 2002), *(iii)* goat kid maternal recognition is based on other vocal features than F0 and the formants (Sèbe *et al*., 2011); or (*iv*) goat kid maternal recognition is based on several features and might be more flexible than previously thought, such that when one main feature is modified, kids focus on other features (Charrier *et al*., 2003).

### Fundamental frequency and formants in maternal recognition

We had predicted that goat fundamental frequency (F0) could be a cue used for individual recognition, because its contour in mother contact calls shows a high Potential for Individual Coding (PIC; start, mean and maximum F0; Briefer & McElligott, 2011b). PIC values greater than 1 indicate that the within-individual variation is lower than the between-individual variation, and therefore that the feature is suitable for individual recognition. Similarly, third and fourth formants’ minimum, mean and maximum values have PIC values > 1 in goat mother calls (Briefer & McElligott, 2011b), making them suitable as well for individual recognition. Because we did not find any effect of F0 or formants modification on maternal recognition, our findings are in line with previous observations suggesting that producing individualised vocalisations does not always result in individual recognition (Carlson *et al*., 2020). Indeed, sometimes, less individualised features might be used by receivers to identify the sender. For example, in pandas (*Ailuropoda melanoleuca*), mean F0 is highly individualised, but females use amplitude modulations instead for recognition of male conspecifics (Charlton *et al*., 2009a). Our results suggest that goat kids might not use F0 and the formants for individual recognition of their mothers, despite these parameters being highly individualised.

We expected that kids would react less to modified than natural vocalisation of their mothers. In order to avoid predators in the wild, it is more adaptive for goat kids to only reply to the vocalisation of their mothers and not reveal their location to unfamiliar individuals or potential predators (Briefer & McElligott, 2011a; Padilla de la Torre *et al*., 2016). In Australian sea lions, pups look and approach the speaker less when F0 of their mothers’ vocalisations have been changed (of once, twice or three times the standard deviation) than when hearing their mothers’ natural vocalisations (Charrier *et al*., 2009). By contrast, fur seals pups have a high tolerance to variation in vocal parameters (Charrier *et al*., 2003); they respond to their mothers’ calls, even when shifted to a degree to which in return their mother can’t recognise them. In our study, goat kids did not differentiate between natural and modified calls of their mothers even when the intra-individual variation was exceeded. This might suggest that goat kids are rather tolerant to variation in parameters, at least those studied here (F0 and formants).

Goat maternal recognition may alternatively rely on duration, amplitude modulation or frequency modulation, which also differ between individuals,, although to a lesser extent than source-filter parameters (Briefer & McElligott, 2011a). In lambs, suppressing amplitude modulation while keeping the natural frequency modulation was found to prevent lambs from identifying their mother’s voice (Sèbe *et al*., 2011). In fur seals, pups can recognise their mother calls despite a suppression of the amplitude modulation, but their recognition ability is impaired by a reversed temporal frequency pattern (Charrier *et al*., 2003). In the same study, fur seal pups could recognise their mother’s call based on the first 25% of the call, but the recognition was impaired if only 10 or 20% of the call was broadcasted, and there was no recognition with only the last 25% of the call (Charrier *et al*., 2003). In goats, the spectral energy distribution was also found to be individualised and could be another parameter on which maternal recognition could be based (Briefer & McElligott, 2011a).

### Effect of the PSOLA based algorithm on the acoustic pattern of goat mothers vocalisations

Despite F0 and formants being theoretically independent of each other (Taylor & Reby, 2010), modifying formants using a PSOLA based algorithm had an impact on the mean, maximum and minimum value of the fundamental frequency, while a modification of fundamental frequency impacted the first, third and fourth formant as well as formants dispersion (see results in Supplementary Material). These results imply that, when measuring the behavioural response to the targeted modified factor, there was still a possibility that subjects’ behaviour would reflect on the unwanted modifications of the other factor. However, even in natural goat vocalisations, F0 and formants values are also correlated to some extent, (Briefer & McElligott, 2011a, Supplementary Material 3). The fundamental nature of the algorithm we used could explain partially these unwanted effects. PSOLA-based algorithm for frequential modifications rely on the precision pitch marks (Rudresh *et al*., 2018) and it has been found to be of insufficient quality for large F0 changes, and particularly very high F0 alterations, by creating imperfections and serious errors (Owsianny, 2019). Considering how goats F0 are naturally quite variable and high, and were positively shifted in the present experiment, the PSOLA algorithm may have produced stimuli where the overall pattern of the call was not preserved, particularly in the second shift treatments where the F0-formants relation was different from the natural contact calls of mothers. Nevertheless, goat kids responded to our shifted modified calls in the same way as natural calls, suggesting that they perceived them as natural calls, and do not rely on F0 and formants, which were all modified to some extent in our playbacks.

## Conclusion

To conclude, our results suggest that, even though they are individualised (Briefer & McElligott, 2011a), source-filter features of the mother’s voice are not used as individual key features in maternal recognition in goats. Further, to our knowledge, our study is the first experimental investigation of the implications of source-filter parameters in offspring recognition of their mothers’ call using a PSOLA-based algorithm. Despite the playback itself influencing kid behaviour, no difference was found between the reaction to the natural vocalisations and reactions to modified ones. It is hence likely that goat kid recognition of mother calls is facilitated by complex relationships between a suite of parameters, rather than individual ones, thus making recognition in varying environmental conditions more robust.

## Supporting information

Scripts

## Acknowledgments

E. Briefer was funded by a Swiss National Science Foundation fellowship. We thank the staff of White Post Farm (http://whitepostfarmcentre.co.uk/) for their help and free access to their animals.

