## Supplementary material for "Goat kid recognition of their mothers’ calls is not impacted by changes in source-filter parameters": Table S1; Table S2; Table S3; Table S4; Table S5; Table S6; Table S7; Table S8

Table S1: Intra-observer reliability for video coding in BORIS Software. 10 videos were randomly selected and coded twice, before comparing each behaviour encoded in both videos using Excel function r². All behaviours had a good intra-observer reliability (min: 0.793; max: 0.992).

| Encoded behaviour | Coefficient of determination (r²) |
| --- | --- |
| Number of calls emitted before playback onset | 0.992 |
| Number of calls emitted during playback broadcast | 0.991 |
| Latency of the first call after playback onset | 0.987 |
| Duration of locomotion before playback onset | 0.953 |
| Duration of locomotion during playback broadcast | 0.901 |
| Latency of the first locomotion after playback onset | 0.868 |
| Duration of looks towards the loudspeaker before playback onset | 0.793 |
| Duration of looks towards the loudspeaker during playback onset | 0.809 |
| Latency of the first looks towards the loudspeaker after playback onset | 0.964 |

*Impact of the PSOLA algorithm on F0 - formants relation*

Shifting the F0 of mother contact calls impacted the formants (LMM; Mean F1, p < 0.01; mean third formant, p < 0.01; mean fourth formant, p < 0.01; formant dispersion, p < 0.01). Further pairwise post-hoc comparisons showed that the natural voice of the mother and F0 Shift1 condition did not differ in their mean first formant, but differed in their mean third formant, mean fourth formant and formant dispersion. Natural voice of the mother and F0 Shift2 condition did not differ in their mean first formant, but differed in their mean third formant, mean fourth formant and formant dispersion. F0Shift1 and F0Shift2 conditions did not differ in their mean first formant, mean third formant, mean fourth formant and formant dispersion. Precise values can be found in Table S2.

Table S2: Relationship between Formant values and F0 measures in mother goat calls in three conditions: natural calls, F0 shift 1 (i.e., F0 positively shifted by one time the standard deviation of the caller), and F0 shift 2 (i.e., F0 shifted by twice the natural standard deviation of the caller). The results of Tukey’s test for each measure in which the model was significantly influenced by the treatment condition are listed (i.e., four out of five measures). Significant terms’ p-values are given in bold.

| **Measure** | **Condition** | **Z/T Ratio** | **P value** |
| --- | --- | --- | --- |
| F1 Mean | F0Shift1 – F0Shift2 | -2.146 | 0.080 |
|  | F0Shift1 - Natural | 0.000 | 1.000 |
|  | F0Shift2 - Natural | 2.146 | 0.080 |
| F3 Mean | F0Shift1 – F0Shift2 | 0.257 | 0.964 |
|  | F0Shift1 - Natural | 2.960 | **0.014** |
|  | F0Shift2 - Natural | 2.702 | 0.272 |
| F4 Mean | F0Shift1 – F0Shift2 | 1.177 | 0.473 |
|  | F0Shift1 - Natural | -2.637 | **0.031** |
|  | F0Shift2 - Natural | -3.814 | **0.001** |
| Formant Dispersion | F0Shift1 – F0Shift2 | 1.153 | 0.488 |
|  | F0Shift1 - Natural | -2.728 | **0.025** |
|  | F0Shift2 - Natural | -3.881 | **0.001** |

Shifting the Formants of mother contact calls impacted the F0 (LMM; mean F0, p < 0.001; maximum F0, p<0.05; and minimum F0, p<0.001). The natural voice of the mother and the Formant shift1 condition differed in their mean F0, max F0 and min F0. The natural voice of the mother and the Formant shift2 condition differed in their mean F0, max F0 and min F0 (p<0.0001). The first formant shift (Shift1 condition) of the voice of the mother and formant shift2 condition did not differ in their mean F0, max F0 and min F0. Precise values can be found in Table S3.

Table S3: Relationship between Formants values and F0 measures in mother goat calls in three conditions: the natural call, the Formant shift 1 (i.e., Formants positively shifted by one time the standard deviation of the caller), and Formant shift 2 (i.e., Formants shifted by twice the natural standard deviation of the caller). The results of Tukey’s test for each measure in which the model was significantly influenced by the treatment condition are listed (i.e., three out of four measures). Significant terms’ p-values are given in bold.

| **Measure** | **Condition** | **Z/T Ratio** | **P value** |
| --- | --- | --- | --- |
| Mean F0 | FormantShift1 – FormantShift2 | 0.830 | 0.687 |
|  | FormantShift1 - Natural | 3.814 | **0.001** |
|  | FormantShift2 - Natural | 2.984 | **0.013** |
| Max F0 | FormantShift1 – FormantShift2 | 0.134 | 0.990 |
|  | FormantShift1 - Natural | 2.599 | **0.034** |
|  | FormantShift2 - Natural | 2.465 | **0.047** |
| Min F0 | FormantShift1 – FormantShift2 | -0.517 | 0.863 |
|  | FormantShift1 - Natural | 7.607 | **<0.0001** |
|  | FormantShift2 - Natural | 8.123 | **<0.0001** |

Table S4: Different model types used throughout the statistical analysis. We checked the residuals of the models graphically for normal distribution and homoscedasticity (simulateResiduals function, package DHARMa, Hartig, 2018). If the assumptions were not met, a logarithmic transformation was used. When the assumptions of normality and homoscedasticity were not met despite a logarithmic transformation, the data were transformed to binary data (superior to the median in the treatment = 1, inferior to the median = 0) and input into Generalized Linear Mixed-Effect Models (GLMM) instead of a LMM.

| **MODEL** | **TYPE OF MODEL** | **TRANSFORMATION (if any)** |
| --- | --- | --- |
| CallRate~ Period+ (1\|Mother/Kid) | LMM |  |
| LocomotionRate~ Period+ (1\|Mother/Kid) | LMM |  |
| LookRate~ Period+ (1\|Mother/Kid) | LMM |  |
| LatencyCall~ Order + DMax_F0 + (1\|Mother/Kid) | GLMM |  |
| LatencyDep~ Order + DMax_F0 + (1\|Mother/Kid) | LMM |  |
| LatencyLook~ Order + DMax_F0 + (1\|Mother/Kid) | LMM | Log+10 |
| CallRateAfter~ Order + DMax_F0 + CallRateBefore + (1\|Mother/Kid) | LMM |  |
| LocomotionRateAfter~ Order + DMax_F0 + LocomotionRateBefore + (1\|Mother/Kid) | LMM |  |
| LookRateAfter~ Order + DMax_F0 + LookRateBefore + (1\|Mother/Kid) | LMM |  |
| LatencyCall~ Order + DMean_F0 + (1\|Mother/Kid) | GLMM |  |
| LatencyDep~ Order + DMean_F0 + (1\|Mother/Kid) | LMM |  |
| LatencyLook~ Order + DMean_F0 + (1\|Mother/Kid) | LMM | Log+10 |
| LocomotionRateAfter~ Order + DMean_F0 + LocomotionRateBefore + (1\|Mother/Kid) | LMM |  |
| LookRateAfter~ Order + DMean_F0 + LookRateBefore + (1\|Mother/Kid) | LMM |  |
| LatencyCall~ Order + DMin_F0 + (1\|Mother/Kid) | GLMM |  |
| LatencyDep~ Order + DMin_F0 + (1\|Mother/Kid) | LMM |  |
| LatencyLook~ Order + DMin_F0 + (1\|Mother/Kid) | LMM | Log+10 |
| CallRateAfter~ Order + DMin_F0 + CallRateBefore + (1\|Mother/Kid) | LMM |  |
| LocomotionRateAfter~ Order + DMin_F0 + LocomotionRateBefore + (1\|Mother/Kid) | LMM |  |
| LookRateAfter~ Order + DMin_F0 + LookRateBefore + (1\|Mother/Kid) | LMM |  |
| LatencyCall~ Order + DRange_F0 + (1\|Mother/Kid) | GLMM |  |
| LatencyDep~ Order + DRange_F0 + (1\|Mother/Kid) | LMM |  |
| LatencyLook~ Order + DRange_F0 + (1\|Mother/Kid) | LMM | Log+10 |
| CallRateAfter~ Order + DRange_F0 + CallRateBefore + (1\|Mother/Kid) | LMM |  |
| LocomotionRateAfter~ Order + DRange_F0 + LocomotionRateBefore + (1\|Mother/Kid) | LMM |  |
| LookRateAfter~ Order + DRange_F0 + LookRateBefore + (1\|Mother/Kid) | LMM |  |
| LatencyCall~ Order + DFDisp + (1\|Mother/Kid) | GLMM |  |
| LatencyDep~ Order + DFDisp + (1\|Mother/Kid) | GLMM |  |
| LatencyLook~ Order + DFDisp + (1\|Mother/Kid) | LMM | Log+10 |
| CallRateAfter~ Order + DFDisp + CallRateBefore + (1\|Mother/Kid) | LMM |  |
| LocomotionRateAfter~ Order + DFDisp + LocomotionRateBefore + (1\|Mother/Kid) | LMM |  |
| LookRateAfter~ Order + DFDisp + LookRateBefore + (1\|Mother/Kid) | LMM |  |
| LatencyCall~ Order + DF1_mean + (1\|Mother/Kid) | GLMM |  |
| LatencyDep~ Order + DF1_mean + (1\|Mother/Kid) | LMM |  |
| LatencyLook~ Order + DF1_mean + (1\|Mother/Kid) | GLMM |  |
| CallRateAfter~ Order + DF1_mean + CallRateBefore + (1\|Mother/Kid) | LMM |  |
| LocomotionRateAfter~ Order + DF1_mean + LocomotionRateBefore + (1\|Mother/Kid) | LMM |  |
| LookRateAfter~ Order + DF1_mean + LookRateBefore + (1\|Mother/Kid) | LMM |  |
| LatencyCall~ Order + DF2_mean + (1\|Mother/Kid) | GLMM |  |
| LatencyDep~ Order + DF2_mean + (1\|Mother/Kid) | LMM |  |
| LatencyLook~ Order + DF2_mean + (1\|Mother/Kid) | LMM | Log+10 |
| CallRateAfter~ Order + DF2_mean + CallRateBefore + (1\|Mother/Kid) | LMM |  |
| LocomotionRateAfter~ Order + DF2_mean + LocomotionRateBefore + (1\|Mother/Kid) | LMM |  |
| LookRateAfter~ Order + DF2_mean + LookRateBefore + (1\|Mother/Kid) | LMM |  |
| LatencyCall~ Order + DF3_mean + (1\|Mother/Kid) | GLMM |  |
| LatencyDep~ Order + DF3_mean + (1\|Mother/Kid) | LMM |  |
| LatencyLook~ Order + DF3_mean + (1\|Mother/Kid) | GLMM |  |
| CallRateAfter~ Order + DF3_mean + CallRateBefore + (1\|Mother/Kid) | LMM |  |
| LocomotionRateAfter~ Order + DF3_mean + LocomotionRateBefore + (1\|Mother/Kid) | LMM |  |
| LookRateAfter~ Order + DF3_mean + LookRateBefore + (1\|Mother/Kid) | LMM |  |
| LatencyCall~ Order + DF4_mean + (1\|Mother/Kid) | GLMM |  |
| LatencyDep~ Order + DF4_mean + (1\|Mother/Kid) | LMM | Log+10 |
| LatencyLook~ Order + DF4_mean + (1\|Mother/Kid) | GLMM |  |
| CallRateAfter~ Order + DF4_mean + CallRateBefore + (1\|Mother/Kid) | LMM |  |
| LocomotionRateAfter~ Order + DF4_mean + LocomotionRateBefore + (1\|Mother/Kid) | LMM |  |
| LookRateAfter~ Order + DF4_mean + LookRateBefore + (1\|Mother/Kid) | LMM |  |
| F1_mean~ Playback + (1\|Mother) | GLMM |  |
| F2_mean~ Playback + (1\|Mother) | LMM |  |
| F3_mean~ Playback + (1\|Mother) | LMM |  |
| F4_mean~ Playback + (1\|Mother) | LMM |  |
| Fdisp~ Playback + (1\|Mother) | LMM |  |
| Mean_F0~ Playback + (1\|Mother) | LMM |  |
| Max_F0~ Playback + (1\|Mother) | LMM |  |
| Min_F0~ Playback + (1\|Mother) | LMM |  |
| Range_F0~ Playback + (1\|Mother) | LMM |  |

Table S5: The order of the treatments affected nine models (LMMs; p value) performed on goat kids’ behaviour in response to the playbacks of their mother’s vocalisations (over three conditions: natural voice of the mother (no shift), a positive shift of F0 within the natural range, and a positive shift of F0 exceeding the natural range). Results were obtained using “drop1” function in the package “stats” on R software. Significant results appear in bold.

|  | Latency of first call | Latency of first locomotion | Latency of fist look towards the speaker | Call Rate | Locomotion  Ratio | Looking rate |
| --- | --- | --- | --- | --- | --- | --- |
| Max F0 | 0.455 | 0.1642 | 0.308 | **0.001** | 0.068 | **0.035** |
| Mean F0 | 0.402 | 0.135 | 0.263 | **0.0006** | **0.041** | **0.025** |
| Min F0 | 0.399 | 0.182 | 0.261 | **0.0009** | **0.042** | **0.014** |
| Range F0 | 0.545 | 0.243 | 0.503 | **0.001** | 0.138 | 0.051 |

Table S6: The behaviour before the playback onset affected all rate behaviours in all 12 models (LMMs; p values) performed on goat kids’ behaviour in response to the playbacks of their mother’s vocalisations (over three conditions: natural voice of the mother (0Hz), a positive shift of F0 the natural range, and a positive shift of F0 exceeding the natural range). Results were obtained using “drop1” function in the package “stats” on R software. Significant results appear in bold.

|  | Call Rate | Locomotion Ratio | Looking ratio |
| --- | --- | --- | --- |
| Max F0 | **0.004** | **0.019** | **0.007** |
| Mean F0 | **0.004** | **0.014** | **0.005** |
| Min F0 | **0.003** | **0.013** | **0.003** |
| Range F0 | **0.002** | **0.034** | **0.005** |

Table S7: The order of the treatments affected all models for locomotion and looking towards the loudspeaker latencies, and for call and locomotion ratios (LMMs; p value) in response to the playbacks of their mother’s vocalisations (over three conditions: natural voice of the mother (no shift), a positive shift of Formants within the natural range, and a positive shift of Formants exceeding the natural range). Results were obtained using “drop1” function in the package “stats” on R software. Significant results appear in bold.

|  | Latency of first call | Latency of first locomotion | Latency of fist look towards the speaker | Call Rate | Locomotion Ratio | Looking ratio |
| --- | --- | --- | --- | --- | --- | --- |
| Mean F1 | 0.691 | **0.018** | **0.032** | **0.037** | **0.001** | 0.285 |
| Mean F2 | 0.647 | **0.019** | **0.005** | **0.037** | **0.001** | 0.278 |
| Mean F3 | 0.609 | **0.016** | **0.024** | **0.037** | **0.001** | 0.262 |
| Mean F4 | 0.405 | **0.001** | **0.013** | **0.037** | **0.0005** | 0.335 |
| Formants Dispersion | 0.373 | **0.004** | **0.002** | **0.036** | **0.0006** | 0.334 |

Table S8: The behaviour before the playback onset affected all rate behaviours in all models performed for call rate and locomotion ratios (LMMs; p values) in response to the playbacks of their mother’s vocalisations (over three conditions: natural voice of the mother (0Hz), a positive shift of Formants within the natural range, and a positive shift of Formants exceeding the natural range). Results were obtained using “drop1” function in the package “stats” on R software. Significant results appear in bold.

|  | Call Rate | Locomotion Ratio | Looking ratio |
| --- | --- | --- | --- |
| Mean F1 | **0.001** | **0.00007** | 0.198 |
| Mean F2 | **0.001** | **0.00008** | 0.201 |
| Mean F3 | **0.001** | **0.00008** | 0.184 |
| Mean F4 | **0.001** | **0.00007** | 0.114 |
| Formants Dispersion | **0.001** | **0.00008** | 0.080 |
