## Supplementary material for "Goat kid recognition of their mothers’ calls is not impacted by changes in source-filter parameters": Scripts

Script 1: Selection of the behaviour rates and ratios affected by the playback (i.e., goats behaviour was significantly different before compared to after the playback onset).

CR=call rate, DR= locomotion ratio, LR= looking ratio

library(lme4)

library (DHARMa)

library (pbkrtest)

library(multcomp)

library(car)

library(MuMIn)

library(lmerTest)

Data_article_CR=read.table("clipboard", h=T, dec = ",")

ModcompCRBvsA<-lmer( CallRate~ Period+ (1|Mother/Kid), REML = F, data = Data_article_CR, na.action=na.omit)

max.simResid<-simulateResiduals ( ModcompCRBvsA, 1000)

plotSimulatedResiduals (max.simResid)

drop1(ModcompCRBvsA,test="Chi")

r.squaredGLMM(ModcompCRBvsA)

Data_article_DR=read.table("clipboard", h=T, dec = ",")

ModcompDRBvsA<-lmer(DepRate~ Period+ (1|Mother/Kid), REML = F, data = Data_article_DR, na.action=na.omit)

max.simResid<-simulateResiduals ( ModcompDRBvsA, 1000)

plotSimulatedResiduals (max.simResid)

drop1(ModcompDRBvsA,test="Chi")

r.squaredGLMM(ModcompDRBvsA)

Data_article_LR=read.table("clipboard", h=T, dec = ",")

ModcompLRBvsA<-lmer(LookRate~ Period+ (1|Mother/Kid), REML = F, data = Data_article_LR, na.action=na.omit)

max.simResid<-simulateResiduals ( ModcompLRBvsA, 1000)

plotSimulatedResiduals (max.simResid)

drop1(ModcompLRBvsA,test="Chi")

r.squaredGLMM(ModcompLRBvsA)

Script 2: Effect of the playback treatment on the responses of the kids for the F0 conditions (natural, F0 Shift1 and F0 Shift2), for each extracted acoustic parameter, the mean acoustic values of the calls of each mother in each treatment were subtracted to the mean value of the natural treatment, to obtain the actual shift (*i.e.,* the natural vocalisation was therefore fixed at zero, and each value given actually refers to that difference, hereafter “playback shift”). One LMM model was built for each selected behaviour, entered as an outcome variable, and for each acoustic parameter measured

CR=call rate, DR= locomotion ratio, LR= looking ratio, CL= Latency of the first call after playback onset, DL= Latency of the first locomotion after playback onset, LL= Latency of the first look after playback onset

F0SelectedRawData=read.table("clipboard",header=TRUE,dec = ",")

F0SelectedBinaryData=read.table("clipboard",header=TRUE,dec = ",")

as.factor(F0SelectedBinaryData$Order)

F0fullmodLC<-lmer((LatencyCall)^1/2~ Order + DRange_F0 + (1|Mother/Kid), REML = F, data = F0SelectedRawData, na.action=na.omit)

max.simResidLC<-simulateResiduals (F0fullmodLC, 1000)

plotSimulatedResiduals (max.simResidLC)

F0fullmodLC<-glmer(LatencyCall~ Order + DRange_F0 + (1|Mother/Kid), family = binomial, data = F0SelectedBinaryData, na.action=na.omit)

F0partmodLC<-glmer(LatencyCall~ Order + (1|Mother/Kid), family = binomial, data = F0SelectedBinaryData, na.action=na.omit)

summary (PBmodcomp(F0fullmodLC, F0partmodLC))

drop1(F0fullmodLC,test="Chi")

r.squaredGLMM(F0fullmodLC)

F0fullmodLD<-lmer(LatencyDep~ Order + DRange_F0 + (1|Mother/Kid), REML = F, data = F0SelectedRawData, na.action=na.omit)

max.simResidLD<-simulateResiduals (F0fullmodLD, 1000)

plotSimulatedResiduals (max.simResidLD)

F0partmodLD<-lmer(LatencyDep~ Order + (1|Mother/Kid), REML = F, data = F0SelectedRawData, na.action=na.omit)

summary (PBmodcomp(F0fullmodLD, F0partmodLD))

drop1(F0fullmodLD,test="Chi")

r.squaredGLMM(F0fullmodLD)

F0fullmodLL<-lmer(log(LatencyLook+10)~ Order + DRange_F0 + (1|Mother/Kid), REML = F, data = F0SelectedRawData, na.action=na.omit)

max.simResidLL<-simulateResiduals (F0fullmodLL, 1000)

plotSimulatedResiduals (max.simResidLL)

F0partmodLL<-lmer(log(LatencyLook+10)~ Order + (1|Mother/Kid), REML = F, data = F0SelectedRawData, na.action=na.omit)

summary (PBmodcomp(F0fullmodLL, F0partmodLL))

drop1(F0fullmodLL,test="Chi")

r.squaredGLMM(F0fullmodLL)

F0fullmodCR<-lmer(CallRateAfter~ Order + DRange_F0 + CallRateBefore + (1|Mother/Kid), REML = F, data = F0SelectedRawData, na.action=na.omit)

max.simResidCR<-simulateResiduals (F0fullmodCR, 1000)

plotSimulatedResiduals (max.simResidCR)

F0partmodCR<-lmer(CallRateAfter~ Order + CallRateBefore + (1|Mother/Kid), REML = F, data = F0SelectedRawData, na.action=na.omit)

summary (PBmodcomp(F0fullmodCR, F0partmodCR))

drop1(F0fullmodCR,test="Chi")

r.squaredGLMM(F0fullmodCR)

F0fullmodDR<-lmer(DepRateAfter~ Order + DRange_F0 + DepRateBefore + (1|Mother/Kid), REML = F, data = F0SelectedRawData, na.action=na.omit)

max.simResidDR<-simulateResiduals (F0fullmodDR, 1000)

plotSimulatedResiduals (max.simResidDR)

F0partmodDR<-lmer(DepRateAfter~ Order + DepRateBefore + (1|Mother/Kid), REML = F, data = F0SelectedRawData, na.action=na.omit)

summary (PBmodcomp(F0fullmodDR, F0partmodDR))

drop1(F0fullmodDR,test="Chi")

r.squaredGLMM(F0fullmodDR)

F0fullmodLR<-lmer(LookRateAfter~ Order + DRange_F0 + LookRateBefore + (1|Mother/Kid), REML = F, data = F0SelectedRawData, na.action=na.omit)

max.simResidLR<-simulateResiduals (F0fullmodLR, 1000)

plotSimulatedResiduals (max.simResidLR)

F0partmodLR<-lmer(LookRateAfter~ Order + LookRateBefore + (1|Mother/Kid), REML = F, data = F0SelectedRawData, na.action=na.omit)

summary (PBmodcomp(F0fullmodLR, F0partmodLR))

drop1(F0fullmodLR,test="Chi")

r.squaredGLMM(F0fullmodLR)

F0fullmodLC<-lmer((LatencyCall)^1/2~ Order + DMean_F0 + (1|Mother/Kid), REML = F, data = F0SelectedRawData, na.action=na.omit)

max.simResidLC<-simulateResiduals (F0fullmodLC, 1000)

plotSimulatedResiduals (max.simResidLC)

F0fullmodLC<-glmer(LatencyCall~ Order + DMean_F0 + (1|Mother/Kid), family = binomial, data = F0SelectedBinaryData, na.action=na.omit)

F0partmodLC<-glmer(LatencyCall~ Order + (1|Mother/Kid), family = binomial, data = F0SelectedBinaryData, na.action=na.omit)

summary (PBmodcomp(F0fullmodLC, F0partmodLC))

drop1(F0fullmodLC,test="Chi")

r.squaredGLMM(F0fullmodLC)

F0fullmodLD<-lmer(LatencyDep~ Order + DMean_F0 + (1|Mother/Kid), REML = F, data = F0SelectedRawData, na.action=na.omit)

max.simResidLD<-simulateResiduals (F0fullmodLD, 1000)

plotSimulatedResiduals (max.simResidLD)

F0partmodLD<-lmer(LatencyDep~ Order + (1|Mother/Kid), REML = F, data = F0SelectedRawData, na.action=na.omit)

summary (PBmodcomp(F0fullmodLD, F0partmodLD))

drop1(F0fullmodLD,test="Chi")

r.squaredGLMM(F0fullmodLD)

F0fullmodLL<-lmer(log(LatencyLook+10)~ Order + DMean_F0 + (1|Mother/Kid), REML = F, data = F0SelectedRawData, na.action=na.omit)

max.simResidLL<-simulateResiduals (F0fullmodLL, 1000)

plotSimulatedResiduals (max.simResidLL)

F0partmodLL<-lmer(log(LatencyLook+10)~ Order + (1|Mother/Kid), REML = F, data = F0SelectedRawData, na.action=na.omit)

summary (PBmodcomp(F0fullmodLL, F0partmodLL))

drop1(F0fullmodLL,test="Chi")

r.squaredGLMM(F0fullmodLL)

F0fullmodCR<-lmer(CallRateAfter~ Order + DMean_F0 + CallRateBefore + (1|Mother/Kid), REML = F, data = F0SelectedRawData, na.action=na.omit)

max.simResidCR<-simulateResiduals (F0fullmodCR, 1000)

plotSimulatedResiduals (max.simResidCR)

F0partmodCR<-lmer(CallRateAfter~ Order + CallRateBefore + (1|Mother/Kid), REML = F, data = F0SelectedRawData, na.action=na.omit)

summary (PBmodcomp(F0fullmodCR, F0partmodCR))

drop1(F0fullmodCR,test="Chi")

r.squaredGLMM(F0fullmodCR)

F0fullmodDR<-lmer(DepRateAfter~ Order + DMean_F0 + DepRateBefore + (1|Mother/Kid), REML = F, data = F0SelectedRawData, na.action=na.omit)

max.simResidDR<-simulateResiduals (F0fullmodDR, 1000)

plotSimulatedResiduals (max.simResidDR)

F0partmodDR<-lmer(DepRateAfter~ Order + DepRateBefore + (1|Mother/Kid), REML = F, data = F0SelectedRawData, na.action=na.omit)

summary (PBmodcomp(F0fullmodDR, F0partmodDR))

r.squaredGLMM(F0fullmodDR)

plot(DepRateAfter~ DMean_F0,F0SelectedRawData,pch=20,ylab= "Locomotion Ratio after playback onset", xlab = "Mean F0 Shift in Hz")

abline(lm(DepRateAfter~ DMean_F0, data = F0SelectedRawData))

conf_interval <- predict(lm(DepRateAfter~ DMean_F0, data = F0SelectedRawData),

newdata = data.frame(DMean_F0 = seq(0, 300, by = 50)),

interval = "confidence", level = 0.95)

lines(seq(0, 300, by = 50), conf_interval[,2], col = "grey", lty = 2)

lines(seq(0, 300, by = 50), conf_interval[,3], col = "grey", lty = 2)

F0fullmodLR<-lmer(LookRateAfter~ Order + DMean_F0 + LookRateBefore + (1|Mother/Kid), REML = F, data = F0SelectedRawData, na.action=na.omit)

max.simResidLR<-simulateResiduals (F0fullmodLR, 1000)

plotSimulatedResiduals (max.simResidLR)

F0partmodLR<-lmer(LookRateAfter~ Order + LookRateBefore + (1|Mother/Kid), REML = F, data = F0SelectedRawData, na.action=na.omit)

summary (PBmodcomp(F0fullmodLR, F0partmodLR))

drop1(F0fullmodLR,test="Chi")

r.squaredGLMM(F0fullmodLR)

F0fullmodLC<-lmer((LatencyCall)^1/2~ Order + DMin_F0 + (1|Mother/Kid), REML = F, data = F0SelectedRawData, na.action=na.omit)

max.simResidLC<-simulateResiduals (F0fullmodLC, 1000)

plotSimulatedResiduals (max.simResidLC)

F0fullmodLC<-glmer(LatencyCall~ Order + DMin_F0 + (1|Mother/Kid), family = binomial, data = F0SelectedBinaryData, na.action=na.omit)

F0partmodLC<-glmer(LatencyCall~ Order + (1|Mother/Kid), family = binomial, data = F0SelectedBinaryData, na.action=na.omit)

summary (PBmodcomp(F0fullmodLC, F0partmodLC))

drop1(F0fullmodLC,test="Chi")

r.squaredGLMM(F0fullmodLC)

F0fullmodLD<-lmer(LatencyDep~ Order + DMin_F0 + (1|Mother/Kid), REML = F, data = F0SelectedRawData, na.action=na.omit)

max.simResidLD<-simulateResiduals (F0fullmodLD, 1000)

plotSimulatedResiduals (max.simResidLD)

F0partmodLD<-lmer(LatencyDep~ Order + (1|Mother/Kid), REML = F, data = F0SelectedRawData, na.action=na.omit)

summary (PBmodcomp(F0fullmodLD, F0partmodLD))

drop1(F0fullmodLD,test="Chi")

r.squaredGLMM(F0fullmodLD)

F0fullmodLL<-lmer(log(LatencyLook+10)~ Order + DMin_F0 + (1|Mother/Kid), REML = F, data = F0SelectedRawData, na.action=na.omit)

max.simResidLL<-simulateResiduals (F0fullmodLL, 1000)

plotSimulatedResiduals (max.simResidLL)

F0partmodLL<-lmer(log(LatencyLook+10)~ Order + (1|Mother/Kid), REML = F, data = F0SelectedRawData, na.action=na.omit)

summary (PBmodcomp(F0fullmodLL, F0partmodLL))

drop1(F0fullmodLL,test="Chi")

r.squaredGLMM(F0fullmodLL)

F0fullmodCR<-lmer(CallRateAfter~ Order + DMin_F0 + CallRateBefore + (1|Mother/Kid), REML = F, data = F0SelectedRawData, na.action=na.omit)

max.simResidCR<-simulateResiduals (F0fullmodCR, 1000)

plotSimulatedResiduals (max.simResidCR)

F0partmodCR<-lmer(CallRateAfter~ Order + CallRateBefore + (1|Mother/Kid), REML = F, data = F0SelectedRawData, na.action=na.omit)

summary (PBmodcomp(F0fullmodCR, F0partmodCR))

drop1(F0fullmodCR,test="Chi")

r.squaredGLMM(F0fullmodCR)

F0fullmodDR<-lmer(DepRateAfter~ Order + DMin_F0 + DepRateBefore + (1|Mother/Kid), REML = F, data = F0SelectedRawData, na.action=na.omit)

max.simResidDR<-simulateResiduals (F0fullmodDR, 1000)

plotSimulatedResiduals (max.simResidDR)

F0partmodDR<-lmer(DepRateAfter~ Order + DepRateBefore + (1|Mother/Kid), REML = F, data = F0SelectedRawData, na.action=na.omit)

summary (PBmodcomp(F0fullmodDR, F0partmodDR))

drop1(F0fullmodDR,test="Chi")

r.squaredGLMM(F0fullmodDR)

plot(DepRateAfter~ DMin_F0,F0SelectedRawData,pch=20,ylab= "Locomotion Ratio after playback onset", xlab = "Min F0 Shift in Hz")

abline(lm(DepRateAfter~ DMin_F0, data = F0SelectedRawData))

conf_interval <- predict(lm(DepRateAfter~ DMin_F0, data = F0SelectedRawData),

newdata = data.frame(DMin_F0 = seq(0, 300, by = 50)),

interval = "confidence", level = 0.95)

lines(seq(0, 300, by = 50), conf_interval[,2], col = "grey", lty = 2)

lines(seq(0, 300, by = 50), conf_interval[,3], col = "grey", lty = 2)

F0fullmodLR<-lmer(LookRateAfter~ Order + DMin_F0 + LookRateBefore + (1|Mother/Kid), REML = F, data = F0SelectedRawData, na.action=na.omit)

max.simResidLR<-simulateResiduals (F0fullmodLR, 1000)

plotSimulatedResiduals (max.simResidLR)

F0aprtmodLR<-lmer(LookRateAfter~ Order + LookRateBefore + (1|Mother/Kid), REML = F, data = F0SelectedRawData, na.action=na.omit)

summary (PBmodcomp(F0fullmodLR, F0partmodLR))

drop1(F0fullmodLR,test="Chi")

r.squaredGLMM(F0fullmodLR)

F0fullmodLC<-lmer((LatencyCall)^1/2~ Order + DMax_F0 + (1|Mother/Kid), REML = F, data = F0SelectedRawData, na.action=na.omit)

max.simResidLC<-simulateResiduals (F0fullmodLC, 1000)

plotSimulatedResiduals (max.simResidLC)

F0fullmodLC<-glmer(LatencyCall~ Order + DMax_F0 + (1|Mother/Kid), family = binomial, data = F0SelectedBinaryData, na.action=na.omit)

F0partmodLC<-glmer(LatencyCall~ Order + (1|Mother/Kid), family = binomial, data = F0SelectedBinaryData, na.action=na.omit)

summary (PBmodcomp(F0fullmodLC, F0partmodLC))

drop1(F0fullmodLC,test="Chi")

r.squaredGLMM(F0fullmodLC)

F0fullmodLD<-lmer(LatencyDep~ Order + DMax_F0 + (1|Mother/Kid), REML = F, data = F0SelectedRawData, na.action=na.omit)

max.simResidLD<-simulateResiduals (F0fullmodLD, 1000)

plotSimulatedResiduals (max.simResidLD)

F0partmodLD<-lmer(LatencyDep~ Order + (1|Mother/Kid), REML = F, data = F0SelectedRawData, na.action=na.omit)

summary (PBmodcomp(F0fullmodLD, F0partmodLD))

drop1(F0fullmodLD,test="Chi")

r.squaredGLMM(F0fullmodLD)

F0fullmodLL<-lmer(log(LatencyLook+10)~ Order + DMax_F0 + (1|Mother/Kid), REML = F, data = F0SelectedRawData, na.action=na.omit)

max.simResidLL<-simulateResiduals (F0fullmodLL, 1000)

plotSimulatedResiduals (max.simResidLL)

F0partmodLL<-lmer(log(LatencyLook+10)~ Order + (1|Mother/Kid), REML = F, data = F0SelectedRawData, na.action=na.omit)

summary (PBmodcomp(F0fullmodLL, F0partmodLL))

drop1(F0fullmodLL,test="Chi")

r.squaredGLMM(F0fullmodLL)

F0fullmodCR<-lmer(CallRateAfter~ Order + DMax_F0 + CallRateBefore + (1|Mother/Kid), REML = F, data = F0SelectedRawData, na.action=na.omit)

max.simResidCR<-simulateResiduals (F0fullmodCR, 1000)

plotSimulatedResiduals (max.simResidCR)

F0partmodCR<-lmer(CallRateAfter~ Order + CallRateBefore + (1|Mother/Kid), REML = F, data = F0SelectedRawData, na.action=na.omit)

summary (PBmodcomp(F0fullmodCR, F0partmodCR))

drop1(F0fullmodCR,test="Chi")

r.squaredGLMM(F0fullmodCR)

F0fullmodDR<-lmer(DepRateAfter~ Order + DMax_F0 + DepRateBefore + (1|Mother/Kid), REML = F, data = F0SelectedRawData, na.action=na.omit)

max.simResidDR<-simulateResiduals (F0fullmodDR, 1000)

plotSimulatedResiduals (max.simResidDR)

F0partmodDR<-lmer(DepRateAfter~ Order + DepRateBefore + (1|Mother/Kid), REML = F, data = F0SelectedRawData, na.action=na.omit)

summary (PBmodcomp(F0fullmodDR, F0partmodDR))

drop1(F0fullmodDR,test="Chi")

r.squaredGLMM(F0fullmodDR)

F0fullmodLR<-lmer(LookRateAfter~ Order + DMax_F0 + LookRateBefore + (1|Mother/Kid), REML = F, data = F0SelectedRawData, na.action=na.omit)

max.simResidLR<-simulateResiduals (F0fullmodLR, 1000)

plotSimulatedResiduals (max.simResidLR)

F0partmodLR<-lmer(LookRateAfter~ Order + LookRateBefore + (1|Mother/Kid), REML = F, data = F0SelectedRawData, na.action=na.omit)

summary (PBmodcomp(F0fullmodLR, F0partmodLR))

drop1(F0fullmodLR,test="Chi")

r.squaredGLMM(F0fullmodLR)

Script 3: Effect of the playback treatment on the responses of the kids for the Formant conditions (natural, Formant Shift1 and Formant Shift2), for each extracted acoustic parameter, the mean acoustic values of the calls of each mother in each treatment were subtracted to the mean value of the natural treatment, to obtain the actual shift (*i.e.,* the natural vocalisation was therefore fixed at zero, and each value given actually refers to that difference, hereafter “playback shift”). One LMM model was built for each selected behaviour, entered as an outcome variable, and for each acoustic parameter measured

CR=call rate, DR= locomotion ratio, LR= looking ratio, CL= Latency of the first call after playback onset, DL= Latency of the first locomotion after playback onset, LL= Latency of the first look after playback onset

FormantsSelectedRawData=read.table("clipboard",header=TRUE,dec = ",")

FormantsSelectedBinaryData=read.table("clipboard",header=TRUE,dec = ",")

as.factor(FormantsSelectedBinaryData$Order)

FormantsfullmodLC<-lmer((LatencyCall)^1/2~ Order + DFDisp + (1|Mother/Kid), REML = F, data = FormantsSelectedRawData, na.action=na.omit)

max.simResidLC<-simulateResiduals (FormantsfullmodLC, 1000)

plotSimulatedResiduals (max.simResidLC)

FormantsfullmodLC<-glmer(LatencyCall~ Order + DFDisp + (1|Mother/Kid), family = binomial, data = FormantsSelectedBinaryData, na.action=na.omit)

FormantspartmodLC<-glmer(LatencyCall~ Order + (1|Mother/Kid), family = binomial, data = FormantsSelectedBinaryData, na.action=na.omit)

summary (PBmodcomp(FormantsfullmodLC, FormantspartmodLC))

drop1(FormantsfullmodLC,test="Chi")

r.squaredGLMM(FormantsfullmodLC)

FormantsfullmodLD<-lmer((LatencyDep)^1/2~ Order + DFDisp + (1|Mother/Kid), REML = F, data = FormantsSelectedRawData, na.action=na.omit)

max.simResidLD<-simulateResiduals (FormantsfullmodLD, 1000)

plotSimulatedResiduals (max.simResidLD)

FormantsfullmodLD<-glmer(LatencyDep~ Order + DFDisp + (1|Mother/Kid), family = binomial, data = FormantsSelectedBinaryData, na.action=na.omit)

FormantspartmodLD<-glmer(LatencyDep~ Order + (1|Mother/Kid), family = binomial, data = FormantsSelectedBinaryData, na.action=na.omit)

summary (PBmodcomp(FormantsfullmodLD, FormantspartmodLD))

drop1(FormantsfullmodLD,test="Chi")

r.squaredGLMM(FormantsfullmodLD)

plot(LatencyDep~DFDisp,FormantsSelectedRawData,pch=20,ylab= "Latency of first locomotion after playback onset (seconds)", xlab = "Shift in Formants Dispersion in Hz")

abline(lm(LatencyDep~DFDisp, data = FormantsSelectedRawData))

conf_interval <- predict(lm(LatencyDep~DFDisp, data = FormantsSelectedRawData),

newdata = data.frame(DFDisp = seq(-60, 400, by = 50)),

interval = "confidence", level = 0.95)

lines(seq(-60, 400, by = 50), conf_interval[,2], col = "grey", lty = 2)

lines(seq(-60, 400, by = 50), conf_interval[,3], col = "grey", lty = 2)

FormantsfullmodLL<-lmer(log(LatencyLook+10)~ Order + DFDisp + (1|Mother/Kid), REML = F, data = FormantsSelectedRawData, na.action=na.omit)

max.simResidLL<-simulateResiduals (FormantsfullmodLL, 1000)

plotSimulatedResiduals (max.simResidLL)

FormantspartmodLL<-lmer(log(LatencyLook+10)~ Order + (1|Mother/Kid), REML = F, data = FormantsSelectedRawData, na.action=na.omit)

summary (PBmodcomp(FormantsfullmodLL, FormantspartmodLL))

drop1(FormantsfullmodLL,test="Chi")

r.squaredGLMM(FormantsfullmodLL)

FormantsfullmodCR<-lmer(CallRateAfter~ Order + DFDisp + CallRateBefore + (1|Mother/Kid), REML = F, data = FormantsSelectedRawData, na.action=na.omit)

max.simResidCR<-simulateResiduals (FormantsfullmodCR, 1000)

plotSimulatedResiduals (max.simResidCR)

FormantspartmodCR<-lmer(CallRateAfter~ Order + CallRateBefore + (1|Mother/Kid), REML = F, data = FormantsSelectedRawData, na.action=na.omit)

summary (PBmodcomp(FormantsfullmodCR, FormantspartmodCR))

drop1(FormantsfullmodCR,test="Chi")

r.squaredGLMM(FormantsfullmodCR)

FormantsfullmodDR<-lmer(DepRateAfter~ Order + DFDisp + DepRateBefore + (1|Mother/Kid), REML = F, data = FormantsSelectedRawData, na.action=na.omit)

max.simResidDR<-simulateResiduals (FormantsfullmodDR, 1000)

plotSimulatedResiduals (max.simResidDR)

FormantspartmodDR<-lmer(DepRateAfter~ Order + DepRateBefore + (1|Mother/Kid), REML = F, data = FormantsSelectedRawData, na.action=na.omit)

summary (PBmodcomp(FormantsfullmodDR, FormantspartmodDR))

drop1(FormantsfullmodDR,test="Chi")

r.squaredGLMM(FormantsfullmodDR)

plot(DepRateAfter~DFDisp,FormantsSelectedRawData,pch=20,ylab= "Locomotion Ratio after playback onset", xlab = "Shift in Formants Dispersion in Hz")

abline(lm(DepRateAfter~DFDisp, data = FormantsSelectedRawData))

conf_interval <- predict(lm(DepRateAfter~DFDisp, data = FormantsSelectedRawData),

newdata = data.frame(DFDisp = seq(-60, 400, by = 50)),

interval = "confidence", level = 0.95)

lines(seq(-60, 400, by = 50), conf_interval[,2], col = "grey", lty = 2)

lines(seq(-60, 400, by = 50), conf_interval[,3], col = "grey", lty = 2)

FormantsfullmodLR<-lmer(LookRateAfter~ Order + DFDisp + LookRateBefore + (1|Mother/Kid), REML = F, data = FormantsSelectedRawData, na.action=na.omit)

max.simResidLR<-simulateResiduals (FormantsfullmodLR, 1000)

plotSimulatedResiduals (max.simResidLR)

FormantspartmodLR<-lmer(LookRateAfter~ Order + LookRateBefore + (1|Mother/Kid), REML = F, data = FormantsSelectedRawData, na.action=na.omit)

summary (PBmodcomp(FormantsfullmodLR, FormantspartmodLR))

drop1(FormantsfullmodLR,test="Chi")

r.squaredGLMM(FormantsfullmodLR)

FormantsfullmodLC<-lmer((LatencyCall)^1/2~ Order + DF4_mean + (1|Mother/Kid), REML = F, data = FormantsSelectedRawData, na.action=na.omit)

max.simResidLC<-simulateResiduals (FormantsfullmodLC, 1000)

plotSimulatedResiduals (max.simResidLC)

FormantsfullmodLC<-glmer(LatencyCall~ Order + DF4_mean + (1|Mother/Kid), family = binomial, data = FormantsSelectedBinaryData, na.action=na.omit)

FormantspartmodLC<-glmer(LatencyCall~ Order + (1|Mother/Kid), family = binomial, data = FormantsSelectedBinaryData, na.action=na.omit)

summary (PBmodcomp(FormantsfullmodLC, FormantspartmodLC))

drop1(FormantsfullmodLC,test="Chi")

r.squaredGLMM(FormantsfullmodLC)

FormantsfullmodLD<-lmer(log(LatencyDep+10)~ Order + DF4_mean + (1|Mother/Kid), REML = F, data = FormantsSelectedRawData, na.action=na.omit)

max.simResidLD<-simulateResiduals (FormantsfullmodLD, 1000)

plotSimulatedResiduals (max.simResidLD)

FormantspartmodLD<-lmer(log(LatencyDep+10)~ Order + (1|Mother/Kid), REML = F, data = FormantsSelectedRawData, na.action=na.omit)

summary (PBmodcomp(FormantsfullmodLD, FormantspartmodLD))

drop1(FormantsfullmodLD,test="Chi")

r.squaredGLMM(FormantsfullmodLD)

FormantsfullmodLL<-lmer((LatencyLook)^1/2~ Order + DF4_mean + (1|Mother/Kid), REML = F, data = FormantsSelectedRawData, na.action=na.omit)

max.simResidLL<-simulateResiduals (FormantsfullmodLL, 1000)

plotSimulatedResiduals (max.simResidLL)

FormantsfullmodLL<-glmer(LatencyLook~ Order + DF4_mean + (1|Mother/Kid), family = binomial, data = FormantsSelectedBinaryData, na.action=na.omit)

FormantspartmodLL<-glmer(LatencyLook~ Order + (1|Mother/Kid), family = binomial, data = FormantsSelectedBinaryData, na.action=na.omit)

summary (PBmodcomp(FormantsfullmodLL, FormantspartmodLL))

drop1(FormantsfullmodLL,test="Chi")

r.squaredGLMM(FormantsfullmodLL)

FormantsfullmodCR<-lmer(CallRateAfter~ Order + DF4_mean + CallRateBefore + (1|Mother/Kid), REML = F, data = FormantsSelectedRawData, na.action=na.omit)

max.simResidCR<-simulateResiduals (FormantsfullmodCR, 1000)

plotSimulatedResiduals (max.simResidCR)

FormantspartmodCR<-lmer(CallRateAfter~ Order + CallRateBefore + (1|Mother/Kid), REML = F, data = FormantsSelectedRawData, na.action=na.omit)

summary (PBmodcomp(FormantsfullmodCR, FormantspartmodCR))

drop1(FormantsfullmodCR,test="Chi")

r.squaredGLMM(FormantsfullmodCR)

FormantsfullmodDR<-lmer(DepRateAfter~ Order + DF4_mean + DepRateBefore + (1|Mother/Kid), REML = F, data = FormantsSelectedRawData, na.action=na.omit)

max.simResidDR<-simulateResiduals (FormantsfullmodDR, 1000)

plotSimulatedResiduals (max.simResidDR)

FormantspartmodDR<-lmer(DepRateAfter~ Order + DepRateBefore + (1|Mother/Kid), REML = F, data = FormantsSelectedRawData, na.action=na.omit)

summary (PBmodcomp(FormantsfullmodDR, FormantspartmodDR))

drop1(FormantsfullmodDR,test="Chi")

r.squaredGLMM(FormantsfullmodDR)

plot(DepRateAfter~DF4_mean,FormantsSelectedRawData,pch=20, ylab= "Locomotion Ratio after playback onset", xlab = "Mean F4 Shift in Hz" )

abline(lm(DepRateAfter~DF4_mean, data = FormantsSelectedRawData))

conf_interval <- predict(lm(DepRateAfter~DF4_mean, data = FormantsSelectedRawData),

newdata = data.frame(DF4_mean= seq(-60, 1500, by = 50)),

interval = "confidence", level = 0.95)

lines(seq(-60, 1500, by = 50), conf_interval[,2], col = "grey", lty = 2)

lines(seq(-60, 1500, by = 50), conf_interval[,3], col = "grey", lty = 2)

FormantsfullmodLR<-lmer(LookRateAfter~ Order + DF4_mean + LookRateBefore + (1|Mother/Kid), REML = F, data = FormantsSelectedRawData, na.action=na.omit)

max.simResidLR<-simulateResiduals (FormantsfullmodLR, 1000)

plotSimulatedResiduals (max.simResidLR)

FormantspartmodLR<-lmer(LookRateAfter~ Order + LookRateBefore + (1|Mother/Kid), REML = F, data = FormantsSelectedRawData, na.action=na.omit)

summary (PBmodcomp(FormantsfullmodLR, FormantspartmodLR))

drop1(FormantsfullmodLR,test="Chi")

r.squaredGLMM(FormantsfullmodLR)

FormantsfullmodLC<-lmer((LatencyCall)^1/2~ Order + DF3_mean + (1|Mother/Kid), REML = F, data = FormantsSelectedRawData, na.action=na.omit)

max.simResidLC<-simulateResiduals (FormantsfullmodLC, 1000)

plotSimulatedResiduals (max.simResidLC)

FormantsfullmodLC<-glmer(LatencyCall~ Order + DF3_mean + (1|Mother/Kid), family = binomial, data = FormantsSelectedBinaryData, na.action=na.omit)

FormantspartmodLC<-glmer(LatencyCall~ Order + (1|Mother/Kid), family = binomial, data = FormantsSelectedBinaryData, na.action=na.omit)

summary (PBmodcomp(FormantsfullmodLC, FormantspartmodLC))

drop1(FormantsfullmodLC,test="Chi")

r.squaredGLMM(FormantsfullmodLC)

FormantsfullmodLD<-lmer(LatencyDep~ Order + DF3_mean + (1|Mother/Kid), REML = F, data = FormantsSelectedRawData, na.action=na.omit)

max.simResidLD<-simulateResiduals (FormantsfullmodLD, 1000)

plotSimulatedResiduals (max.simResidLD)

FormantspartmodLD<-lmer(LatencyDep~ Order + (1|Mother/Kid), REML = F, data = FormantsSelectedRawData, na.action=na.omit)

summary (PBmodcomp(FormantsfullmodLD, FormantspartmodLD))

drop1(FormantsfullmodLD,test="Chi")

r.squaredGLMM(FormantsfullmodLD)

FormantsfullmodLL<-lmer((LatencyLook)^1/2~ Order + DF3_mean + (1|Mother/Kid), REML = F, data = FormantsSelectedRawData, na.action=na.omit)

max.simResidLL<-simulateResiduals (FormantsfullmodLL, 1000)

plotSimulatedResiduals (max.simResidLL)

FormantsfullmodLL<-glmer(LatencyLook~ Order + DF3_mean + (1|Mother/Kid), family = binomial, data = FormantsSelectedBinaryData, na.action=na.omit)

FormantspartmodLL<-glmer(LatencyLook~ Order + (1|Mother/Kid), family = binomial, data = FormantsSelectedBinaryData, na.action=na.omit)

summary (PBmodcomp(FormantsfullmodLL, FormantspartmodLL))

drop1(FormantsfullmodLL,test="Chi")

r.squaredGLMM(FormantsfullmodLL)

FormantsfullmodCR<-lmer(CallRateAfter~ Order + DF3_mean + CallRateBefore + (1|Mother/Kid), REML = F, data = FormantsSelectedRawData, na.action=na.omit)

max.simResidCR<-simulateResiduals (FormantsfullmodCR, 1000)

plotSimulatedResiduals (max.simResidCR)

FormantspartmodCR<-lmer(CallRateAfter~ Order + CallRateBefore + (1|Mother/Kid), REML = F, data = FormantsSelectedRawData, na.action=na.omit)

summary (PBmodcomp(FormantsfullmodCR, FormantspartmodCR))

drop1(FormantsfullmodCR,test="Chi")

r.squaredGLMM(FormantsfullmodCR)

FormantsfullmodDR<-lmer(DepRateAfter~ Order + DF3_mean + DepRateBefore + (1|Mother/Kid), REML = F, data = FormantsSelectedRawData, na.action=na.omit)

max.simResidDR<-simulateResiduals (FormantsfullmodDR, 1000)

plotSimulatedResiduals (max.simResidDR)

FormantspartmodDR<-lmer(DepRateAfter~ Order + DepRateBefore + (1|Mother/Kid), REML = F, data = FormantsSelectedRawData, na.action=na.omit)

summary (PBmodcomp(FormantsfullmodDR, FormantspartmodDR))

drop1(FormantsfullmodDR,test="Chi")

r.squaredGLMM(FormantsfullmodDR)

FormantsfullmodLR<-lmer(LookRateAfter~ Order + DF3_mean + LookRateBefore + (1|Mother/Kid), REML = F, data = FormantsSelectedRawData, na.action=na.omit)

max.simResidLR<-simulateResiduals (FormantsfullmodLR, 1000)

plotSimulatedResiduals (max.simResidLR)

FormantspartmodLR<-lmer(LookRateAfter~ Order + LookRateBefore + (1|Mother/Kid), REML = F, data = FormantsSelectedRawData, na.action=na.omit)

summary (PBmodcomp(FormantsfullmodLR, FormantspartmodLR))

drop1(FormantsfullmodLR,test="Chi")

r.squaredGLMM(FormantsfullmodLR)

FormantsfullmodLC<-lmer((LatencyCall)^1/2~ Order + DF2_mean + (1|Mother/Kid), REML = F, data = FormantsSelectedRawData, na.action=na.omit)

max.simResidLC<-simulateResiduals (FormantsfullmodLC, 1000)

plotSimulatedResiduals (max.simResidLC)

FormantsfullmodLC<-glmer(LatencyCall~ Order + DF2_mean + (1|Mother/Kid), family = binomial, data = FormantsSelectedBinaryData, na.action=na.omit)

FormantspartmodLC<-glmer(LatencyCall~ Order + (1|Mother/Kid), family = binomial, data = FormantsSelectedBinaryData, na.action=na.omit)

summary (PBmodcomp(FormantsfullmodLC, FormantspartmodLC))

drop1(FormantsfullmodLC,test="Chi")

r.squaredGLMM(FormantsfullmodLC)

FormantsfullmodLD<-lmer(LatencyDep~ Order + DF2_mean + (1|Mother/Kid), REML = F, data = FormantsSelectedRawData, na.action=na.omit)

max.simResidLD<-simulateResiduals (FormantsfullmodLD, 1000)

plotSimulatedResiduals (max.simResidLD)

FormantspartmodLD<-lmer(LatencyDep~ Order + (1|Mother/Kid), REML = F, data = FormantsSelectedRawData, na.action=na.omit)

summary (PBmodcomp(FormantsfullmodLD, FormantspartmodLD))

drop1(FormantsfullmodLD,test="Chi")

r.squaredGLMM(FormantsfullmodLD)

FormantsfullmodLL<-lmer(log(LatencyLook+10)~ Order + DF2_mean + (1|Mother/Kid), REML = F, data = FormantsSelectedRawData, na.action=na.omit)

max.simResidLL<-simulateResiduals (FormantsfullmodLL, 1000)

plotSimulatedResiduals (max.simResidLL)

FormantspartmodLL<-lmer(log(LatencyLook+10)~ Order + (1|Mother/Kid), REML = F, data = FormantsSelectedRawData, na.action=na.omit)

summary (PBmodcomp(FormantsfullmodLL, FormantspartmodLL))

drop1(FormantsfullmodLL,test="Chi")

r.squaredGLMM(FormantsfullmodLL)

FormantsfullmodCR<-lmer(CallRateAfter~ Order + DF2_mean + CallRateBefore + (1|Mother/Kid), REML = F, data = FormantsSelectedRawData, na.action=na.omit)

max.simResidCR<-simulateResiduals (FormantsfullmodCR, 1000)

plotSimulatedResiduals (max.simResidCR)

FormantspartmodCR<-lmer(CallRateAfter~ Order + CallRateBefore + (1|Mother/Kid), REML = F, data = FormantsSelectedRawData, na.action=na.omit)

summary (PBmodcomp(FormantsfullmodCR, FormantspartmodCR))

drop1(FormantsfullmodCR,test="Chi")

r.squaredGLMM(FormantsfullmodCR)

FormantsfullmodDR<-lmer(DepRateAfter~ Order + DF2_mean + DepRateBefore + (1|Mother/Kid), REML = F, data = FormantsSelectedRawData, na.action=na.omit)

max.simResidDR<-simulateResiduals (FormantsfullmodDR, 1000)

plotSimulatedResiduals (max.simResidDR)

FormantspartmodDR<-lmer(DepRateAfter~ Order + DepRateBefore + (1|Mother/Kid), REML = F, data = FormantsSelectedRawData, na.action=na.omit)

summary (PBmodcomp(FormantsfullmodDR, FormantspartmodDR))

drop1(FormantsfullmodDR,test="Chi")

r.squaredGLMM(FormantsfullmodDR)

FormantsfullmodLR<-lmer(LookRateAfter~ Order + DF2_mean + LookRateBefore + (1|Mother/Kid), REML = F, data = FormantsSelectedRawData, na.action=na.omit)

max.simResidLR<-simulateResiduals (FormantsfullmodLR, 1000)

plotSimulatedResiduals (max.simResidLR)

FormantspartmodLR<-lmer(LookRateAfter~ Order + LookRateBefore + (1|Mother/Kid), REML = F, data = FormantsSelectedRawData, na.action=na.omit)

summary (PBmodcomp(FormantsfullmodLR, FormantspartmodLR))

drop1(FormantsfullmodLR,test="Chi")

r.squaredGLMM(FormantsfullmodLR)

FormantsfullmodLC<-lmer((LatencyCall)^1/2~ Order + DF1_mean + (1|Mother/Kid), REML = F, data = FormantsSelectedRawData, na.action=na.omit)

max.simResidLC<-simulateResiduals (FormantsfullmodLC, 1000)

plotSimulatedResiduals (max.simResidLC)

FormantsfullmodLC<-glmer(LatencyCall~ Order + DF1_mean + (1|Mother/Kid), family = binomial, data = FormantsSelectedBinaryData, na.action=na.omit)

FormantspartmodLC<-glmer(LatencyCall~ Order + (1|Mother/Kid), family = binomial, data = FormantsSelectedBinaryData, na.action=na.omit)

summary (PBmodcomp(FormantsfullmodLC, FormantspartmodLC))

drop1(FormantsfullmodLC,test="Chi")

r.squaredGLMM(FormantsfullmodLC)

FormantsfullmodLD<-lmer(LatencyDep~ Order + DF1_mean + (1|Mother/Kid), REML = F, data = FormantsSelectedRawData, na.action=na.omit)

max.simResidLD<-simulateResiduals (FormantsfullmodLD, 1000)

plotSimulatedResiduals (max.simResidLD)

FormantspartmodLD<-lmer(LatencyDep~ Order + (1|Mother/Kid), REML = F, data = FormantsSelectedRawData, na.action=na.omit)

summary (PBmodcomp(FormantsfullmodLD, FormantspartmodLD))

drop1(FormantsfullmodLD,test="Chi")

r.squaredGLMM(FormantsfullmodLD)

FormantsfullmodLL<-lmer((LatencyLook)^1/2~ Order + DF1_mean + (1|Mother/Kid), REML = F, data = FormantsSelectedRawData, na.action=na.omit)

max.simResidLL<-simulateResiduals (FormantsfullmodLL, 1000)

plotSimulatedResiduals (max.simResidLL)

FormantsfullmodLL<-glmer(LatencyLook~ Order + DF1_mean + (1|Mother/Kid), family = binomial, data = FormantsSelectedBinaryData, na.action=na.omit)

FormantspartmodLL<-glmer(LatencyLook~ Order + (1|Mother/Kid), family = binomial, data = FormantsSelectedBinaryData, na.action=na.omit)

summary (PBmodcomp(FormantsfullmodLL, FormantspartmodLL))

drop1(FormantsfullmodLL,test="Chi")

r.squaredGLMM(FormantsfullmodLL)

FormantsfullmodCR<-lmer(CallRateAfter~ Order + DF1_mean + CallRateBefore + (1|Mother/Kid), REML = F, data = FormantsSelectedRawData, na.action=na.omit)

max.simResidCR<-simulateResiduals (FormantsfullmodCR, 1000)

plotSimulatedResiduals (max.simResidCR)

FormantspartmodCR<-lmer(CallRateAfter~ Order + CallRateBefore + (1|Mother/Kid), REML = F, data = FormantsSelectedRawData, na.action=na.omit)

summary (PBmodcomp(FormantsfullmodCR, FormantspartmodCR))

drop1(FormantsfullmodCR,test="Chi")

r.squaredGLMM(FormantsfullmodCR)

FormantsfullmodDR<-lmer(DepRateAfter~ Order + DF1_mean + DepRateBefore + (1|Mother/Kid), REML = F, data = FormantsSelectedRawData, na.action=na.omit)

max.simResidDR<-simulateResiduals (FormantsfullmodDR, 1000)

plotSimulatedResiduals (max.simResidDR)

FormantspartmodDR<-lmer(DepRateAfter~ Order + DepRateBefore + (1|Mother/Kid), REML = F, data = FormantsSelectedRawData, na.action=na.omit)

summary (PBmodcomp(FormantsfullmodDR, FormantspartmodDR))

drop1(FormantsfullmodDR,test="Chi")

r.squaredGLMM(FormantsfullmodDR)

FormantsfullmodLR<-lmer(LookRateAfter~ Order + DF1_mean + LookRateBefore + (1|Mother/Kid), REML = F, data = FormantsSelectedRawData, na.action=na.omit)

max.simResidLR<-simulateResiduals (FormantsfullmodLR, 1000)

plotSimulatedResiduals (max.simResidLR)

FormantspartmodLR<-lmer(LookRateAfter~ Order + LookRateBefore + (1|Mother/Kid), REML = F, data = FormantsSelectedRawData, na.action=na.omit)

summary (PBmodcomp(FormantsfullmodLR, FormantspartmodLR))

drop1(FormantsfullmodLR,test="Chi")

r.squaredGLMM(FormantsfullmodLR)

Script 4: impact of modifying F0 on formants

F0Control=read.table("clipboard",header=TRUE,dec = ",")

F0ControlBinary=read.table("clipboard",header=TRUE,dec = ",")

as.factor(F0ControlBinary$Order)

F0fullmodCtrlF1mean<-lmer((F1_mean)^1/2~ Playback + (1|Mother), REML = F, data = F0Control, na.action=na.omit)

max.simResidCtrlF1mean<-simulateResiduals (F0fullmodCtrlF1mean, 1000)

plotSimulatedResiduals (max.simResidCtrlF1mean)

F0fullmodCtrlF1mean<-glmer(F1_mean~ Playback + (1|Mother), family = binomial, data = F0ControlBinary, na.action=na.omit)

F0partmodCtrlF1mean<-glmer(F1_mean~ (1|Mother), family = binomial, data = F0ControlBinary, na.action=na.omit)

summary (PBmodcomp(F0fullmodCtrlF1mean, F0partmodCtrlF1mean))

drop1(F0fullmodCtrlF1mean,test="Chi")

r.squaredGLMM(F0fullmodCtrlF1mean)

F0fullmodCtrlF2mean<-lmer(F2_mean~ Playback + (1|Mother), REML = F, data = F0Control, na.action=na.omit)

max.simResidCtrlF2mean<-simulateResiduals (F0fullmodCtrlF2mean, 1000)

plotSimulatedResiduals (max.simResidCtrlF2mean)

F0partmodCtrlF2mean<-lmer(F2_mean~ (1|Mother), REML = F, data = F0Control, na.action=na.omit)

summary (PBmodcomp(F0fullmodCtrlF2mean, F0partmodCtrlF2mean))

drop1(F0fullmodCtrlF2mean,test="Chi")

r.squaredGLMM(F0fullmodCtrlF2mean)

F0fullmodCtrlF3mean<-lmer(F3_mean~ Playback + (1|Mother), REML = F, data = F0Control, na.action=na.omit)

max.simResidCtrlF3mean<-simulateResiduals (F0fullmodCtrlF3mean, 1000)

plotSimulatedResiduals (max.simResidCtrlF3mean)

F0partmodCtrlF3mean<-lmer(F3_mean~ (1|Mother), REML = F, data = F0Control, na.action=na.omit)

summary (PBmodcomp(F0fullmodCtrlF3mean, F0partmodCtrlF3mean))

drop1(F0fullmodCtrlF3mean,test="Chi")

r.squaredGLMM(F0fullmodCtrlF3mean)

F0fullmodCtrlF4mean<-lmer(F4_mean~ Playback + (1|Mother), REML = F, data = F0Control, na.action=na.omit)

max.simResidCtrlF4mean<-simulateResiduals (F0fullmodCtrlF4mean, 1000)

plotSimulatedResiduals (max.simResidCtrlF4mean)

F0partmodCtrlF4mean<-lmer(F4_mean~ (1|Mother), REML = F, data = F0Control, na.action=na.omit)

summary (PBmodcomp(F0fullmodCtrlF4mean, F0partmodCtrlF4mean))

drop1(F0fullmodCtrlF4mean,test="Chi")

r.squaredGLMM(F0fullmodCtrlF4mean)

F0fullmodCtrlFDisp<-lmer(Fdisp~ Playback + (1|Mother), REML = F, data = F0Control, na.action=na.omit)

max.simResidCtrlFDisp<-simulateResiduals (F0fullmodCtrlFDisp, 1000)

plotSimulatedResiduals (max.simResidCtrlFDisp)

F0partmodCtrlFDisp<-lmer(Fdisp~ (1|Mother), REML = F, data = F0Control, na.action=na.omit)

summary (PBmodcomp(F0fullmodCtrlFDisp, F0partmodCtrlFDisp))

drop1(F0fullmodCtrlFDisp,test="Chi")

r.squaredGLMM(F0fullmodCtrlFDisp)

library(emmeans)

emmeans(F0fullmodCtrlF1mean, list(pairwise ~ Playback), adjust = "tukey")

emmeans(F0fullmodCtrlFDisp, list(pairwise ~ Playback), adjust = "tukey")

emmeans(F0fullmodCtrlF3mean, list(pairwise ~ Playback), adjust = "tukey")

emmeans(F0fullmodCtrlF4mean, list(pairwise ~ Playback), adjust = "tukey")

FormantControl=read.table("clipboard",header=TRUE,dec = ",")

F0ControlBinary=read.table("clipboard",header=TRUE,dec = ",")

as.factor(F0ControlBinary$Order)

FormantfullmodCtrlMean_F0<-lmer(Mean_F0~ Playback + (1|Mother), REML = F, data = FormantControl, na.action=na.omit)

max.simResidCtrlMean_F0<-simulateResiduals (FormantfullmodCtrlMean_F0, 1000)

plotSimulatedResiduals (max.simResidCtrlMean_F0)

FormantpartmodCtrlMean_F0<-lmer(Mean_F0~(1|Mother), REML = F, data = FormantControl, na.action=na.omit)

summary (PBmodcomp(FormantfullmodCtrlMean_F0, FormantpartmodCtrlMean_F0))

drop1(FormantfullmodCtrlMean_F0,test="Chi")

r.squaredGLMM(FormantfullmodCtrlMean_F0)

FormantfullmodCtrlMax_F0<-lmer(Max_F0~ Playback + (1|Mother), REML = F, data = FormantControl, na.action=na.omit)

max.simResidCtrlMax_F0<-simulateResiduals (FormantfullmodCtrlMax_F0, 1000)

plotSimulatedResiduals (max.simResidCtrlMax_F0)

FormantpartmodCtrlMax_F0<-lmer(Max_F0~(1|Mother), REML = F, data = FormantControl, na.action=na.omit)

summary (PBmodcomp(FormantfullmodCtrlMax_F0, FormantpartmodCtrlMax_F0))

drop1(FormantfullmodCtrlMax_F0,test="Chi")

r.squaredGLMM(FormantfullmodCtrlMax_F0)

FormantfullmodCtrlMin_F0<-lmer(Min_F0~ Playback + (1|Mother), REML = F, data = FormantControl, na.action=na.omit)

max.simResidCtrlMin_F0<-simulateResiduals (FormantfullmodCtrlMin_F0, 1000)

plotSimulatedResiduals (max.simResidCtrlMin_F0)

FormantpartmodCtrlMin_F0<-lmer(Min_F0~(1|Mother), REML = F, data = FormantControl, na.action=na.omit)

summary (PBmodcomp(FormantfullmodCtrlMin_F0, FormantpartmodCtrlMin_F0))

drop1(FormantfullmodCtrlMin_F0,test="Chi")

r.squaredGLMM(FormantfullmodCtrlMin_F0)

FormantfullmodCtrlRange_F0<-lmer(Range_F0~ Playback + (1|Mother), REML = F, data = FormantControl, na.action=na.omit)

max.simResidCtrlRange_F0<-simulateResiduals (FormantfullmodCtrlRange_F0, 1000)

plotSimulatedResiduals (max.simResidCtrlRange_F0)

FormantpartmodCtrlRange_F0<-lmer(Range_F0~(1|Mother), REML = F, data = FormantControl, na.action=na.omit)

summary (PBmodcomp(FormantfullmodCtrlRange_F0, FormantpartmodCtrlRange_F0))

drop1(FormantfullmodCtrlRange_F0,test="Chi")

r.squaredGLMM(FormantfullmodCtrlRange_F0)

emmeans(FormantfullmodCtrlMean_F0, list(pairwise ~ Playback), adjust = "tukey")

emmeans(FormantfullmodCtrlMin_F0, list(pairwise ~ Playback), adjust = "tukey")

emmeans(FormantfullmodCtrlMax_F0, list(pairwise ~ Playback), adjust = "tukey")

emmeans(FormantfullmodCtrlRange_F0, list(pairwise ~ Playback), adjust = "tukey")
